# Data-driven causal analysis of observational time series in ecology

**DOI:** 10.1101/2020.08.03.233692

**Authors:** Alex E Yuan, Wenying Shou

## Abstract

Complex ecosystems are challenging to understand as they often defy manipulative experiments for practical or ethical reasons. In response, several fields have developed parallel approaches to infer causal relations from observational time series. Yet these methods are easy to misunderstand and often controversial. Here, we provide an accessible and critical review of three statistical causal inference approaches popular in ecological time series analysis: pairwise correlation, Granger causality, and state space reconstruction. For each, we ask what a method tests for, what causal statement it might imply, and when it could lead us astray. We devise new ways of visualizing key concepts, describe some novel pathologies of causal inference methods, and point out how so-called “model-free” causality tests are not assumption-free. We hope that our synthesis will facilitate thoughtful application of causal inference approaches and encourage explicit statements of assumptions.

## Introduction

Ecological communities perform important activities, from facilitating digestion in the human gut to driving the biogeochemical cycles of elements on the earth. Communities are often highly complex, with many species engaging in diverse interactions. To control communities, it helps to know whether the abundance of a species might change as a result of a perturbation in, for instance, the abundance of another species or the availability of a nutrient.

Ideally, biologists discover such causal relations from manipulative experiments. However, manipulative experiments can be infeasible or inappropriate: Natural ecosystems may not offer enough replicates for comprehensive manipulative experiments, and perturbations can be impractical at large scales and may have unanticipated negative consequences. On the other hand, there exists an ever-growing plethora of population dynamic time series which are observational in nature (i.e. without intentional perturbations). The idea that one might wrest reasonably accurate causal predictions from these time series is tantalizing.

Causal inference can become more straightforward if one already knows, or is willing to assume, a model that captures key aspects of the underlying process. For example, the Lotka-Volterra model popular in mathematical ecology assumes that species interact in a pairwise fashion, that the fitness effects from different interactions are additive, and that all pairwise interactions can be represented by a single equation form where parameters can vary to reflect signs and strengths of fitness effects. By fitting such a model to time series of species abundances and environmental factors, one can predict, for instance, which species interact or how a community might respond to certain perturbations [1, 3, 2]. However, the Lotka-Volterra equations often fail to describe complex ecosystems and chemically-mediated interactions [55, 54, 6].

When our understanding is insufficient to support knowledge-based modeling, how might we formulate causal hypotheses? A large and rapidly growing literature of advanced methods attempts to infer causal relations from time series data without using a mechanistic model. Such methods are sometimes called “model-free” [52], although they may rely on *statistical* models. Some of these methods are equation-free (e.g. examining probability distributions), while others deploy equations that are usually not interpreted mechanistically (e.g. structural equation models). Causal inference methods work either implicitly by predicting the effects of new perturbations [75, 77] or explicitly by proposing causal structures [81, 74, 73, 72, 71, 78, 76].

Here we focus on three model-free approaches that have been commonly used to make causal claims in ecology research: pairwise correlation, Granger causality, and state space reconstruction. For each, we ask (1) what information does the method give us, (2) what causal statement might that information imply, and (3) when might the method lead us astray?

We found that answering these seemingly basic questions was surprisingly challenging for several reasons. First, modern causal inference approaches have intellectual roots in several communities including philosophy, statistics, econometrics, and chaos theory, which sometimes use different words for the same idea, and the same word for different ideas. The word causality itself is a prime example: Many philosophers (and scientists) would say that *X* causes *Y* if an intervention upon *X* would result in a change in *Y* [30, 16]. Granger’s original works instead defined causality to be about how important the history of *X* is in predicting *Y* [17, 26], and in the nonlinear dynamics field, causality is sometimes used to mean that the trajectories of *X* and *Y* have certain shared geometric or topological properties [14]. Such language, while unproblematic when confined to a single community, can nevertheless obscure important differences between methods from different communities. A second challenge is that in methodological articles, key assumptions are sometimes hidden in algorithmic details, or simply not mentioned. Finally, some methods deal with nuanced or advanced mathematical ideas that can be difficult even for those with quantitative training. Given these challenges, it is no surprise that research using causal inference from observational time series has been highly controversial, with an abundance of “letters to the editor”, often followed by impassioned dialogue [20, 24, 79, 90, 25].

We strive to balance precision and readability. To accomplish this, we devised new ways to visualize key concepts. We also provide refreshers and discussions of mathematical notions in the Appendices. Lastly, we compare all methods to a common definition of causality that is useful to experimental scientists. Our goals are to inform practitioners who wish to make or evaluate causal hypotheses using temporal data, to facilitate communication across different fields interested in time series causal inference, and to encourage explicit statements of methodological assumptions and caveats.

## Dependence, correlation, and causality

### Causality

In this article, we use the definition of “causality” that is common in statistics and intuitive to scientists: *X* has a causal effect on *Y* (“*X* causes *Y* “ or “*X* is a causer; *Y* is a causee” or “*X* is a cause; *Y* is an effect”) if some externally applied perturbation of *X* can result in a perturbation in *Y* (Figure 1A). We say that *X* and *Y* are *causally related* if *X* causes *Y, Y* causes *X*, or some other variable (“common cause”, “confounder”) causes both. Otherwise, *X* and *Y* are *causally unrelated*. Additionally, one can talk about direct versus indirect causality (Figure 1B; see legend for definitions). Systems for which one can correctly reveal all direct causal relationships between all pairs of variables are called causally identifiable, and various theoretical identifiability conditions are known [81, 8, 80]. However, empirical time series often do not contain enough information for easy causal identifiability [81, 73].

**Figure 1:**
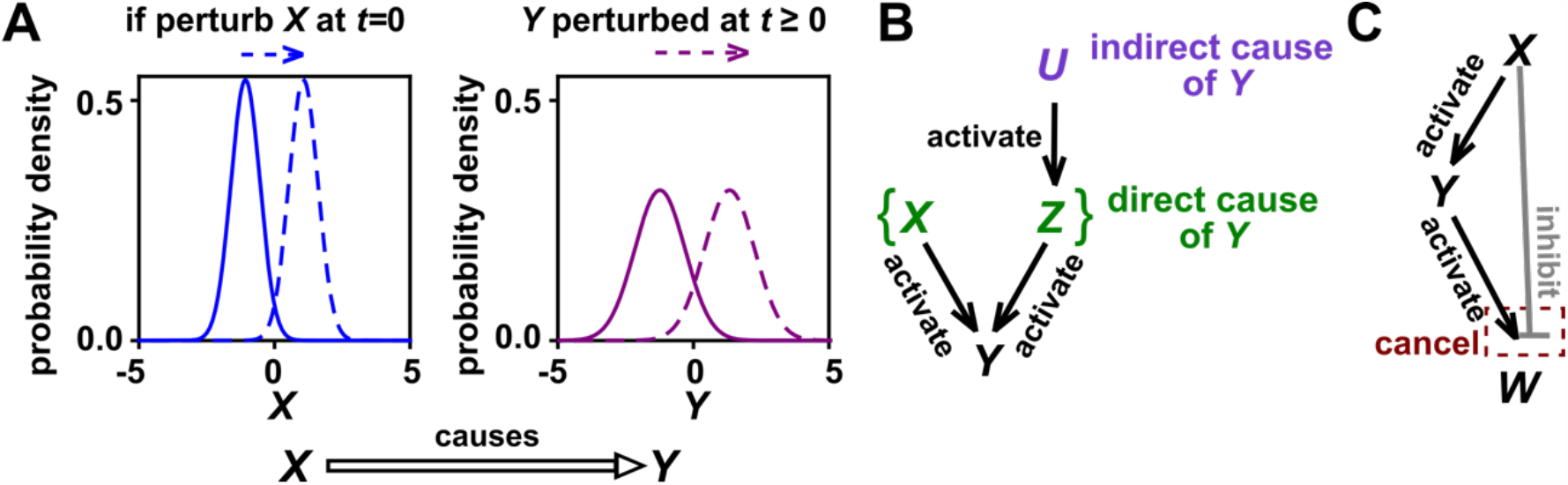
Causality. (**A**) Definition. If a perturbation in *X* results in a change in future (or current) *Y*, then *X* causes *Y*. This definition does not require that *any* perturbation in *X* will perturb *Y*. For example, if the effect of *X* on *Y* has saturated, then a further increase in *X* will not affect *Y*. To embody probabilistic thinking (e.g. drunk driving increases the chance of accidents) [16], *X* and *Y* are depicted as histograms. Perturbing *X* can perturb the current value of *Y* if, for example, *X* and *Y* describe the same quantity or if *X* and *Y* are linked by a conservation law (e.g. conservation of energy). (**B**) Direct versus indirect causality. The direct causers of *Y* are given by the minimal set of variables such that once the entire set is fixed, no other variables can cause *Y*. Here, three players *X, Z*, and *U* activate *Y*. The set {*X, Z*} constitutes the direct causers of *Y* (*Y*’s “Markovian parents” [29, 16]), since if we fix both *X* and *Z*, then *Y* becomes independent of *U*. If a causer is not direct, we say that it is indirect. Whether a causer is direct or indirect can depend on the scope of included variables. For example, suppose that yeast releases acetate, and acetate inhibits the growth of bacteria. If acetate is not in our scope, then yeast density has a direct causal effect on bacterial density. Conversely, if acetate is included in our scope, then acetate (but not yeast) is the direct causer of bacterial density since fixing acetate concentration would fix bacterial growth regardless of yeast density. (**C**) Direct causality does not always imply causality. Here, {*X, Y*} are the direct causers of *W*. While *X* directly causes *W, X* does not cause *W* because its direct effect and indirect effect (through *Y*) cancel out. Also note that causality is not always transitive: *X* causes *Y* and *Y* causes *W*, but *X* does not cause *W*. In this article, causality is represented by a hollow arrow, while activation and inhibition are represented by filled arrows and blunt-head arrows, respectively.

### Correlation versus dependence

The adage “correlation is not causality” is well-known to the point of being cliché [5, 10, 9, 28]. Yet, to dismiss correlative evidence altogether seems too extreme. To make use of correlative evidence without being reckless, it helps to distinguish between the terms “correlation” and “dependence”. When applied to ecological time series, the term “correlation” is often used to mean some statistic that quantifies the similarity between two observed time series [70, 10]. Examples include Pearson’s correlation coefficient and local similarity [33]. In contrast, statistical dependence is a hypothesis about the probability distributions that produced those time series, and has close connections to causality.

Dependence has a precise definition in statistics. Dependence is most easily described for two binary events. For instance, if the incidence of vision loss is higher among diabetics than among the general population, then vision loss and diabetes are statistically dependent. In general, events *A* and *B* are dependent if across many independent trials (e.g. patients), the probability that *A* occurs given that *B* has occurred (e.g. incidence of vision loss among diabetics only) is different from the background probability that *A* occurs (e.g. background incidence of vision loss). *A* and *B* are independent if they are not dependent. The concept of dependence is readily generalized from pairs of binary events to pairs of numerical variables (Appendix 1.1), and then again to pairs of vectors, such as time series (Appendix 1.3, Figure S7).

Dependence is connected to causation by the widely accepted “Common Cause Principle”: *if two variables are dependent, then they are causally related (one causes the other, or both share a common cause)* [8, 71, 80, 44]. Note however that if one mistakenly introduces selection bias, then two independent variables can appear to be dependent (Figure S6). The closely related property of conditional dependence (Appendix 1.3), i.e. whether two variables are dependent after statistically controlling for (i.e. “fixing”) certain other variables, can be even more causally informative. In fact, when conditional dependencies (and conditional independencies) are known, it is sometimes possible to infer most or all of the direct causal relationships at play, even without manipulative experiments or temporal information. Many of the algorithms that accomplish this rely on two technical but often reasonable assumptions: the “causal Markov condition”, which allows one to infer causal information from conditional *dependence*, and the “causal faithfulness condition”, which allows one to infer causal information from conditional *independence* [8, 73, 80].

In sum, whereas a correlation is a statistical description of data, statistical dependence is a hypothesis about the relationship between the underlying probability distributions. Dependence is in turn linked to causality. Below, we discuss how to use correlation to detect dependence in time series.

### Testing for dependence between time series using surrogate data

Despite its scientific usefulness, dependence between time series can be treacherous to test for. This is because time series are often autocorrelated (e.g. what occurs today influences what occurs tomorrow), so that a single pair of time series contains information from only a single trial. If one has many trials that are independent and free of systematic differences (e.g. ≥ 20 as in laboratory microcosm experiments), the task is relatively easy: One can test whether species *X* and *Y* are statistically dependent by comparing the correlation between *X* and *Y* abundance series from the same trial with those between *X* and *Y* abundance series from different trials (Appendix 1.4). However, a large trial number is generally a luxury and often only one trial is available. In such cases, attempting to discern whether two time series are statistically dependent is like attempting to divine whether diabetes and vision loss are dependent with only a single patient (i.e. we have an “*n*-of-one problem”). As one possible remedy, parametric tests that account for autocorrelation can be used to evaluate the statistical significance of a Pearson correlation coefficient [89], although these make multiple technical assumptions that are difficult to verify [88, 23].

In practice, the *n*-of-one problem is often addressed by a technique called surrogate data testing. Specifically, one computes some measure of correlation between two time series *X* and *Y*. Next, one uses a computer to simulate replicates of *Y* that might have been obtained if *X* and *Y* were independent (see below). Each simulated replicate is called a “surrogate” *Y*. Finally, one computes the correlation between *X* and each surrogate *Y*. The *p*-value (the probability of observing the original correlation or a stronger one under the null hypothesis that *X* and *Y* are independent) is then determined by counting how many of the surrogate *Y* s produce a stronger correlation than the real *Y*. For example, if we produced 19 surrogates and found the real correlation to be stronger than all 19 surrogate correlations, then we would write down a *p*-value of 1*/*(1 + 19) = 0.05. Ideally, if two time series are not causally related, then we should register a a *p*-value of 0.05 (or less) in only 5% of cases.

Several procedures are commonly used for generating surrogate *Y* data, each corresponding to an assumption about how the original *Y* time series was generated. One popular procedure is to shuffle the values of *Y* in time [33, 56, 57, 58]. This procedure, often called permutation, assumes that all time points of *Y* are independent of each other. A rejection of this null hypothesis (i.e. a low *p*-value) often provides little evidence of dependence because rarely is this assumption met. For example, for independent time series in Figure 1B-C, this test returns *p* < 0.05 at rates of 30 ∼ 92%, much higher than 5%. Nevertheless, the permutation test sneaks into many applied works, perhaps because it is the default option in some popular software packages. Another procedure for generating surrogate *Y* data has been called phase randomization. It first uses the Fourier transform to represent a time series as a sum of sine waves, then randomly shifts each of the component sine waves in time, and finally sums the phase-shifted components [23, 60] (Figure S8). This procedure assumes that *Y* is Gaussian, linear, and stationary [60, 59], where “Gaussian” means that any subsequence of *Y* follows a multivariate Gaussian distribution, “stationary” means that this distribution does not change over time, and “linear” means that future *Y* values depend linearly on past *Y* values and past random events (“process noise”, Figure 7A). Indeed, this test performed well (with a false positive rate of 4%) when *Y* satisfied the three assumptions (Figure 2C), and relatively poorly when the stationarity assumption was violated (with a false positive rate of 21%; Figure 2B). Other surrogate data procedures include time shifting [60] and the twin method [65]. Some surrogate-based independence tests have been shown to perform reasonably well even when exact theoretical requirements are unmet or unknown [65, 34], but a more comprehensive benchmarking effort is needed.

**Figure 2:**
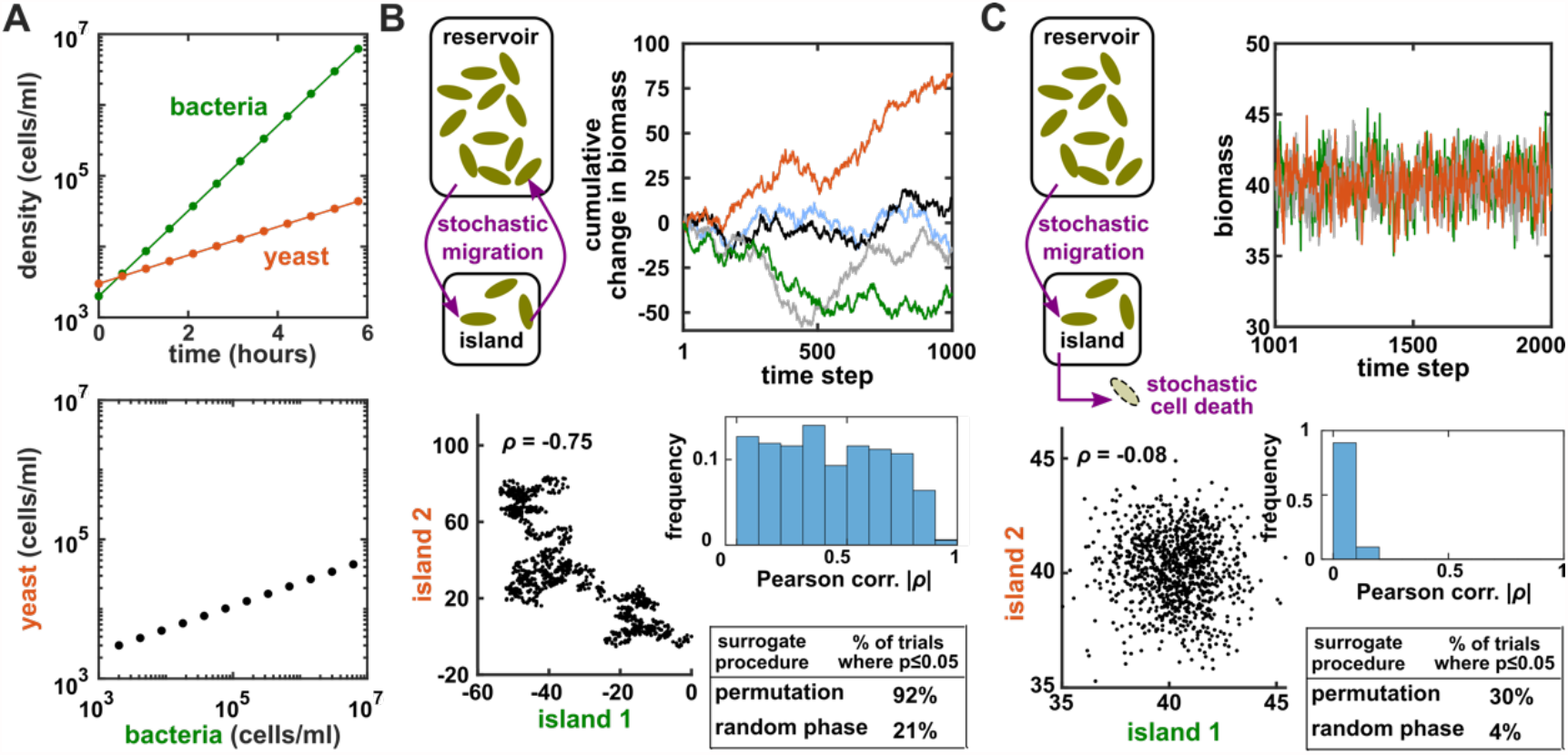
Two independent temporal processes can appear significantly correlated. (**A**) Densities of independent yeast and bacteria cultures growing exponentially are correlated. (**B, C**) Correlation between time series of two independent island populations can appear significant if inappropriate tests are used. (**B**) In an island, individuals stochastically migrate to and from a “reservoir” (a random walk). At each time step, the net change in island biomass is drawn from a standard normal distribution (mean = 0; standard deviation = 1 biomass unit). (**C**) A population receives cells through migration and loses cells to death. Population size has reached an equilibrium. In both cases, we plotted example time series (upper right), a scatterplot comparing two island populations (lower left), the histogram of Pearson correlation coefficient strength (blue shading), and the fraction of simulations in which the correlation was deemed significant (*p* ≤ 0.05) by surrogate data tests using either permutation or phase randomization (see main text). Ideally, the proportion of correlations that are significant (false positives) should not exceed 5%. The strength of correlation is weaker in (C) compared to (B), yet still often significant according to the permutation test. See Supplementary Methods for more details.

In sum, surrogate data allow a researcher to use an observed correlation statistic to test for dependence under some assumption about the data-generating process. Dependence indicates the presence of a causal relationship, and conditional dependence can sometimes even indicate the direction [80, 73, 85] (Figure S5). Below we consider Granger causality and convergent cross mapping, two methods which explicitly take advantage of temporal information to infer the direction of causality.

## Granger causality: intuition, pitfalls, and implementations

### Intuition and formal definitions

In simple language, *X* is said to Granger-cause *Y* if a collection of time series containing all historical measurements predicts *Y*’s future behavior better than a similar collection that excludes the history of *X*. An important consequence of this definition is that Granger causality excludes indirect causes, as illustrated in Figure 3A. In practice, whether a causal relationship is direct or indirect depends on which variables are observed (e.g. in Figure 3A, if *Y* were not observed, then *X* would “directly” and Granger-cause *Z*).

**Figure 3:**
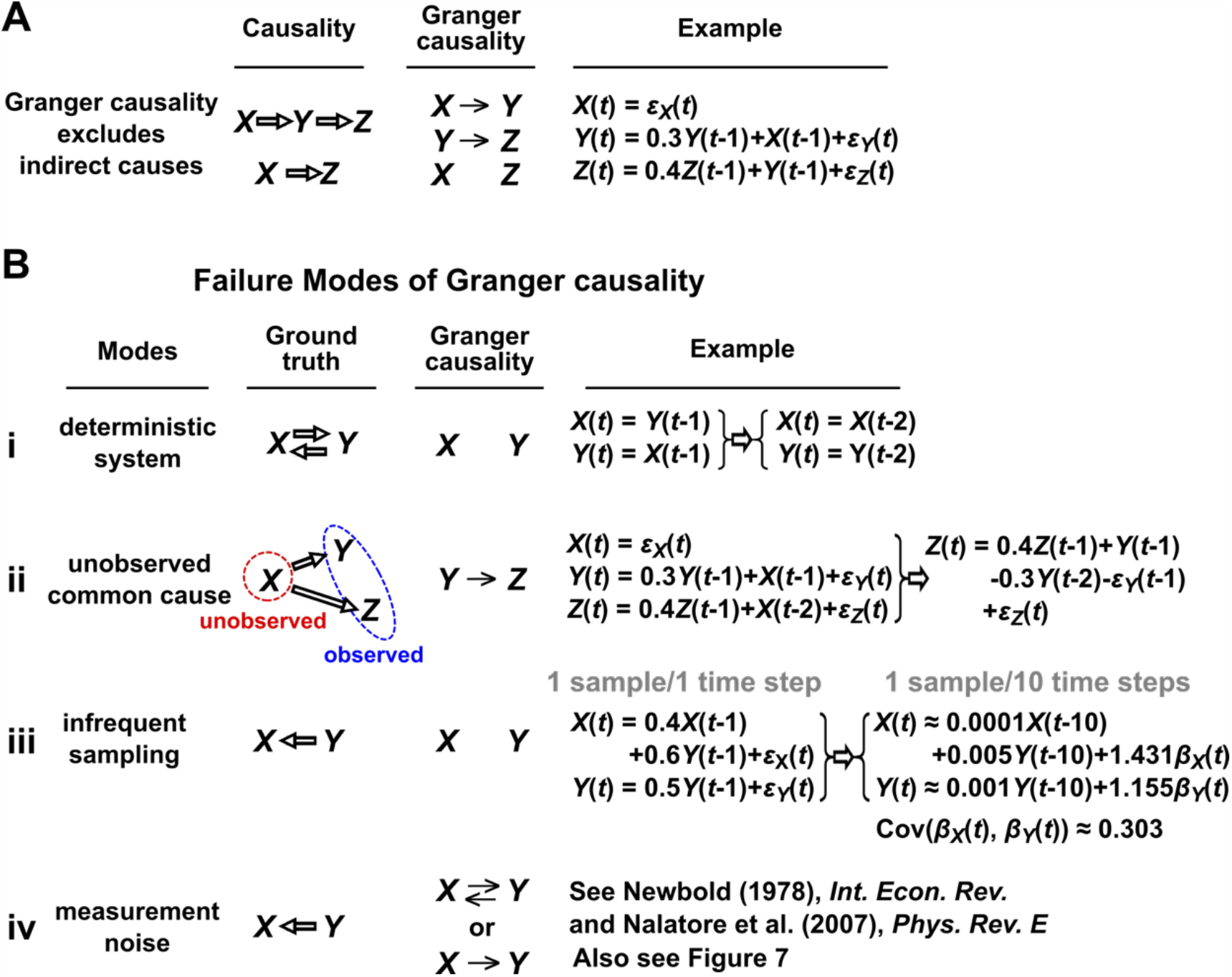
Causality versus Granger causality. (**A**) Granger causality is designed to reveal direct causes, not indirect causes. Although *X* causes *Z, X* does not Granger-cause *Z* because with the history of *Y* available, the history of *X* no longer adds value for predicting *Z*. In this example, we can also see that Granger causality is not transitive: *X* Granger-causes *Y* and *Y* Granger-causes *Z*, but *X* does not Granger-cause *Z*. (**B**) Failure modes of Granger causality when inferring direct causality. (**i**) False negative due to lack of stochasticity. *X* and *Y* mutually and deterministically cause one another through a copy operation [91, 8]: *X*(*t*) copies *Y* (*t* − 1) and vice versa. Since *X*(*t* − 2) already contains sufficient information to know *X*(*t*) exactly, the history of *Y* cannot improve prediction of *X*, and so *Y* does not Granger-cause *X*. By symmetry, *X* does not Granger-cause *Y*. (**ii**) False positive due to unobserved common cause. *X* causes *Y* with a delay of 1, and causes *Z* with a delay of 2. We only observe *Y* and *Z*. Since *Y* receives the same “information” before *Z*, the history of *Y* helps to predict *Z*, and thus *Y* Granger-causes *Z*, resulting in a false positive. (**iii**) Infrequent sampling induces false negatives. Note that in the equation of *X*, the ratio of signal (the coefficient of causer *Y*) to noise (the coefficient of process noise or *“*) is much smaller when sampling is infrequent (compare right with left). The quantitative relationship is derived in Appendix 1.10. (**iv**) Measurement noise can lead Granger causality to suffer both false positives and false negatives. *ϵ*_*X*_, *ϵ*_*Y*_, *ϵ*_*Z*_, *β*_*X*_, and *β*_*Y*_ represent process noise and are normal random variables with mean of 0 and variance of 1. *ϵ*_*X*_, *ϵ*_*Y*_, and *ϵ*_*Z*_ are independent of one another. Coefficients are chosen to ensure stationarity, except in (**ii**).

Granger causality has many related but nonequivalent quantitative incarnations in the literature, including several that were proposed by Granger himself [17, 26]. Box 1 presents two definitions: one based on a linear regression which we call “linear Granger causality” [4, 7, 35, 28] and a second, more general, definition which we call “general Granger causality” (also sometimes called nonlinear Granger causality) [26, 38, 39, 36, 37, 34].

#### Box 1

**Granger causality**

1. **Linear Granger causality:** Under linear Granger causality, *X* Granger-causes *Y* if including the history of *X* in a linear autoregressive model (Eq 1) allows for a better prediction of future *Y* than not including the history of *X* (i.e. setting all *α*_*k*_ coefficients to zero). By “linear autoregressive model”, we mean that the future value of variable *Y* is modeled as a linear combination of historical values of *X* and *Y* and all other observed variables that might help predict *Y* “ … ”:

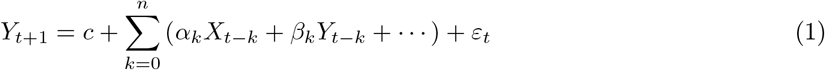 Here, *t* is the time index, *k* = 0, 1, …,*n* is a time lag index, *c* is a constant, coefficients such as *α*_*k*_ and *β*_*k*_ represent the strength of contributions from the respective terms, and *ε*_*t*_ represents independent and identically-distributed (IID, 1.2) process noise (Figure 7A).
2. **General Granger causality [26]:** Let *X*_*t*_ *Y*_*t*_, and *Z*_*t*_ be series of random variables indexed by time *t. X* Granger-causes *Y* with respect to the information set {*X*_*t*_, *Y*_*t*_, *Z*_*t*_} if:

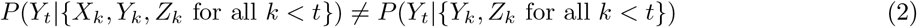

for at least one possible realization of the *X, Y*, and *Z* series. Here, *P*(*Y*_*t*_|𝒮) is the probability of *Y*_*t*_ conditional on the variable set 𝒮. Note that *Z*_*k*_ in Eq. 2 may include multiple variables and thus plays the same role as “ … ” in Eq. 1.

### Failure modes

We discuss four conceptually or practically important instances where Granger causality can fail as an indicator of direct causality (Figure 3B). These pathologies, applicable to both linear and general Granger causality, can be understood intuitively. First, if a system has deterministic dynamics (Appendix 1.9), then Granger causality may fail to detect causal relations (Figure 3Bi). We do not expect population dynamics to be completely deterministic. Nevertheless, when dynamics have a low degree of randomness (i.e. are almost deterministic), Granger causality may suffer low sensitivity [8] and other methods have been developed to address exactly this situation [5]. Second, Granger causality may erroneously assign a direct causal relation between a pair of variables that have an unobserved common cause (Figure 3Bii). Third, when sampling frequency is low (“subsampling”), the causal signal may become too low to detect (Figure 3Biii), although recent techniques in combination with non-Gaussian noise terms may reduce problems associated with subsampling [82, 83]. Lastly, when measurements are noisy due to for instance instrument uncertainty (Figure 3Biv), Granger causality can assign false interactions and also fail to detect true causality [46], although some progress has been made on this front [47].

### Practical testing for linear and general Granger causality

One might still attempt to infer Granger causality despite the above caveats, especially if they can be largely avoided. Linear Granger causality, popular in microbiome studies [4, 7, 35, 28], uses standard parametric tests: if any of the *α*_*k*_ terms in Eq.1 is nonzero, then *X* linear Granger-causes *Y*. Parametric tests are computationally inexpensive and available in multiple free and well-documented software packages [43, 27]. These tests assume that time series are “covariance-stationary”, which means that certain statistical properties of the series are time-independent [27] (see Appendix 1.8), and can fail when this assumption is violated [41, 40, 42]. Additionally, when linear Granger causality is applied to nonlinear systems, it can produce wrong causal inferences, even if none of the hazards from Figure 3B are at play [66]. One can assess whether the linear model (Eq. 1) is a reasonable approximation, for instance by checking whether the model residuals *ε*_*t*_ are uncorrelated across time [48] (as is assumed by Eq. 1).

Tests for general Granger causality often use surrogate data null models, with the same procedures and caveats discussed previously. In these tests, it is common to estimate a statistic known as transfer entropy [86]. Roughly, the transfer entropy from *X* to *Y* is the extent to which the entropy (a measurement of uncertainty) of *Y*’s future is reduced when we account for (specifically, condition on) the past of *X* [92, 64, 87, 34]. Thus, testing for general Granger causality can be accomplished by testing the significance of transfer entropy using surrogate data techniques [87, 34]. Several implementations of such tests are freely available (e.g. [87, 32, 31]).

Granger causality methods face challenges when datasets have a large number of variables (e.g. in microbial ecology). In this case, the summation in Eq. 1 will contain a large number of terms, and so a regression procedure may assign a small importance to each of the terms, but fail to assign a “significant” importance to many true interactions [71, 72]. To handle systems with many variables, one can impose the assumption that only a small number of causal links exist [4, 28]. This is sometimes called sparse regression or regularization. Additionally, under certain technical assumptions, it is possible to use a series of logical rules to remove unnecessary terms in a purely data-driven way. These rules and their associated assumptions are formalized in “constraint-based causal discovery” algorithms [8, 73] (Appendix 1.5). The development of new causal discovery algorithms, and their application to time series, is a very active area of research [72, 71, 78, 84].

## State space reconstruction (SSR): intuition, pitfalls, and implementations

The term “state space reconstruction” (SSR) refers to a broad swath of techniques for prediction, inference, and estimation in time series analysis [69, 67, 68, 5, 15]. In this article, when we use the term SSR, we refer only to SSR methods for causality detection. These methods enjoy widespread use in empirical ecology [49, 50, 63, 62, 61]. SSR methods are intended to complement Granger causality: Whereas Granger causality has trouble with deterministic dynamics (Figure 3B), the SSR approach is explicitly designed for systems that are primarily deterministic [5]. Since SSR is less intuitive than correlation or Granger causality, we introduce it with an example rather than a definition.

### Visualizing SSR causal inference

Consider the deterministic dynamical system in Figure 4. Here, *Z* is causally driven by *Y* (and *X*), but not by *W* or *V*. We can make a vector out of the current value *Z*(*t*) and two past values *Z*(*t* − *τ*) and *Z*(*t* − 2*τ*) (Figure 4B, red dots), where *τ* is the time delay and [*Z*(*t*),*Z*(*t* − *τ*),*Z*(*t* − 2*τ*)] is called a “delay vector”. The delay vector can be represented as a single point in the 3-dimensional *Z* “delay space” (Figure 4C, red dot). We then shade each point of the *Z* delay space according to the contemporaneous value of its causer *Y* (*t*). Since in this example each point of the *Z* delay space corresponds to one and only one *Y* (*t*) value, we call this a “delay map” from *Z* to *Y*. Notice that the *Y* (*t*) gradient in this plot looks gradual in the sense that if two time points are nearby in the delay space of *Z*, then their corresponding *Y* (*t*) shades are also similar. This property is called “continuity” (Figure S12). We provide additional details on continuity, and the more stringent criterion of smoothness, in Appendix 1.11. Overall, there is a continuous map from the *Z* delay space to *Y*, or more concisely, a “continuous delay map” from *Z* to *Y*. A similar continuous delay map also exists from *Z* to its other causer *X*. On the other hand, if we shade the delay space of *Z* by *W* or *V* (neither of which causes *Z*), we do not get a continuous delay map (Figure 4D-E).

**Figure 4:**
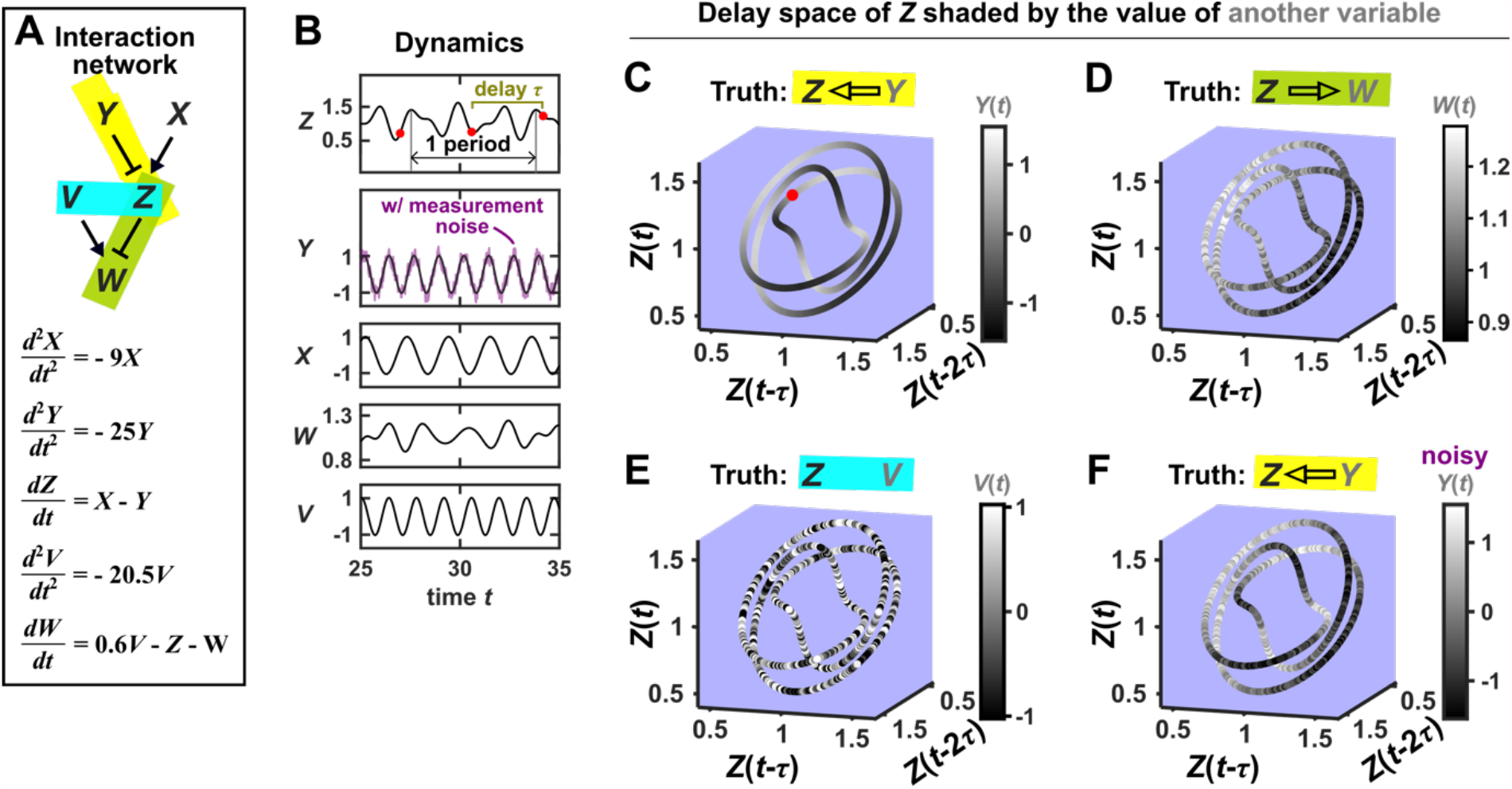
SSR causal inference looks for a continuous map from the delay space of a causee to the causer, and becomes more difficult in the presence of noise. (**A**) A 5-variable toy (linear) system. Filled arrows and blunt head arrows represent activation and inhibition, respectively. (**B**) Time series. The delay vector [*Z*(*t*),*Z*(*t* − *τ*),*Z*(*t* − 2*τ*)] (shown as three red dots) can be represented as a single point in the 3-dimensional *Z* delay space (**C**, red dot). (**C**) We then shade each point in the *Z* delay space by its corresponding contemporaneous value of *Y* (*t*) (without measurement noise). The shading is continuous (i.e. gradual transitions in shade), and note that *Y* causes *Z* (correctly, in this case). (**D**) When we repeat this procedure, but now shade the *Z* delay space by *W* (*t*), the shading is bumpy, and note that *W* does not cause *Z* (even though *Z* causes *W*). (**E**) Shading the delay space of *Z* by the causally unrelated *V* also gives a bumpy result. (**F**) Dynamics as in (**C**), but now with noisy measurements of *Y* (purple in **B**). The shading is no longer gradual. Thus with noisy data, causal inference is more difficult. See Supplementary Methods for more details.

In this example, there is a continuous delay map from a causee to a causer, but not the other way around, and also no continuous delay map between causally unrelated variables. If this behavior reflects a broader principle, then perhaps continuous delay maps can be used to infer the presence and direction of causation. Is there in fact a broader principle?

In fact, there is a sort of broader principle, but it may not be fully satisfying for causal inference. The principle is a classic theorem due to Floris Takens [19]. Very roughly, Takens’s theorem can be interpreted for our situation as follows: If *X* and *Y* are variables in a deterministic dynamical system, and if *X* influences *Y*, and if several other technical requirements are met, then we are almost guaranteed to find a continuous delay map from *Y* to *X*. We walk through these details with visual examples in Appendix 1.13. In principle, this phenomenon is mostly insensitive to the choice of delay *τ* and the length of delay vector (see Appendix 1.12 for practical considerations in delay vector construction). Takens’s theorem and later results [22] form the theoretical basis of SSR techniques. However, Takens’s theorem does *not* state that a continuous delay map implies a causal relationship.

In sum, SSR techniques attempt to detect a smooth delay map (or a related feature) between two variables and use this to infer the presence and direction of causation [5, 45, 14]: A smooth delay map from *Y* to *X* is taken as an indication that *X* causes *Y*. However, Takens’s theorem leaves open the possibility of a smooth delay map without the corresponding causation. Indeed, we now illustrate scenarios where a causal relationship and a continuous delay map do not coincide.

### SSR failure modes

Figure 5 illustrates four failure modes of SSR. In the first failure mode, which we refer to as “nonreverting continuous dynamics” (top row of Figure 5; Appendix 1.14), a continuous map arises from the delay space of *X* to *Z* because a continuous map can be found from the delay space of *X* to time (“nonreverting”) and from time to *Z* (“continuous”). This pathology leads to false causal inference and may explain apparently causal results in some early works where SSR methods were applied to time series with a clear trend. We are not aware of statistical tests for this problem, but Clark et al. [18] recommend shading points in the delay space with their corresponding time to visually check for a time trend. In the second failure mode [5] (Figure 5, second row), one variable drives another variable in such a way that the dynamics of the two variables are synchronized. Consequently, although the true causal relationship is unidirectional, bidirectional causality is inferred. Although the “prediction lag test” (Figure 6B right panel) can sometimes alleviate this problem [11, 12], it is not foolproof as we demonstrate in Appendix 1.15. In the third failure mode (Figure 5 third row; [12]), *X* and *Z* both oscillate and *X*’s period is an integer multiple of *Z*’s period. In this case, *Z* is inferred to cause *X* even though they are causally unrelated. In the fourth failure mode (Figure 5, bottom row), SSR gives a false negative error due to “pathological symmetry”, although this may be rare in practice.

**Figure 5:**
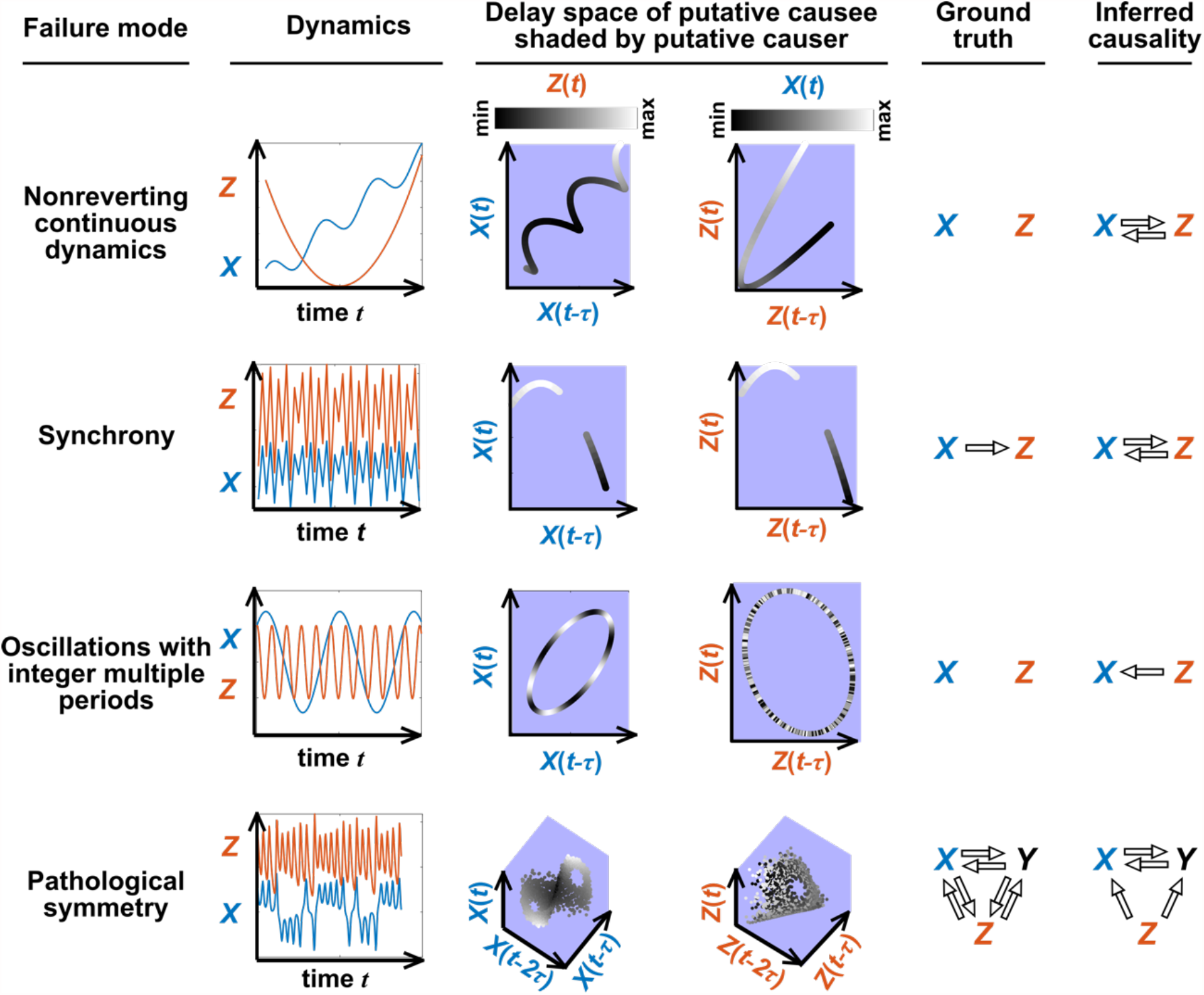
Failure modes associated with state space reconstruction. **Top row**: Nonreverting continuous dynamics may lead one to infer causality where there is none. This example consists of two time series: a wavy linear increase and a parabolic trajectory. Although they are causally unrelated, we can find continuous delay maps between them. This is because there is (i) a continuous map from the delay vector [*X*(*t*),*X*(*t* − *τ*)] to *t* (*X* is “nonreverting”), and (ii) a continuous map from *t* to *Z* (*Z* is “continuous”), and thus there is a continuous delay map from *X* to *Z* (“nonreverting continuous dynamics”). Thus, one falsely infers that *Z* causes *X*, and with similar reasoning that *X* causes *Z*. **Second row**: *X* drives *Z* such that their dynamics are “synchronzied”, and consequently, we find a continuous delay map also from *X* to *Z* even though *Z* does not drive *X*. Note that the extent of synchronization is not always apparent from inspecting equations (e.g. Figure 12 of [13]) or dynamics (Figure S16). **Third row**: *X* oscillates at a period that is 5 times the oscillatory period of *Y*. There is a continuous delay map from *X* to *Z* even through *X* and *Z* are causally unrelated. Note that true causality sometimes also induces oscillations with integer multiples periods (e.g. in Figure 4, the period of *Z* is 3 times the period of *X*). **Bottom row**: In the classic chaotic Lorenz attractor, the *Z* variable is caused by *X* and *Y*, but we do not see a continuous map from the delay space of *Z* to either *X* (shown) or *Y* (not shown). This is because, as mentioned earlier, satisfying the conditions in Takens’s theorem makes a continuous mapping likely but not guaranteed (Appendix 1.13). Here, *Z* is an example of this lack of guarantee [21] due to a symmetry in the system (see “Background definitions for causation in dynamic systems” in the Supplementary Information of [5]).

**Figure 6:**
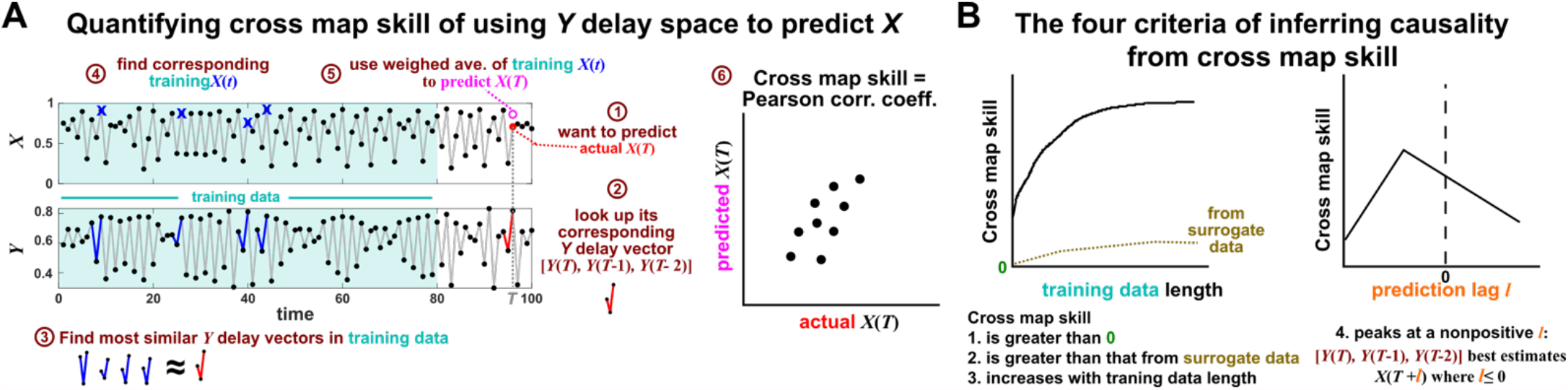
Illustration of the convergent cross mapping (CCM) procedure for testing whether *X* causes *Y*. (**A**) Quantifying cross map skill. Consider the point *X*(*T*) denoted by the red dot (“actual *X*(*T*)” in ➀), which we want to predict from *Y* delay vectors. We first look up the contemporaneous *Y* delay vector ([*Y* (*T*),*Y* (*T* − 1),*Y* (*T* − 2)]) (➁, red dynamics), and identify times within our training data when delay vectors of *Y* were the most similar to our red delay vector (➂, blue segments). We then look up their contemporaneous values of *X* (➃, blue crosses), and use their weighted average to predict *X*(*T*) (➄, open magenta circle; weights are given as equations S2 and S3 in the supplement of [5]). We repeat this procedure for many choices of *T* and calculate the Pearson correlation coefficient between the actual *X*(*T*) and predicted *X*(*T*) (➅). This correlation is called the “cross map skill”. While other measures of cross map skill, such as mean squared error, may also be used [5], our choice follows the convention of [5]. (**B**) Four criteria for inferring causality from the cross map skill. Data points in **(A)** are marked by dots and connecting lines are visual aids.

### Convergent cross mapping: Detecting SSR causal signals from real data

Despite the caveats above, one might still attempt to use SSR to hypothesize causal relations. In this case, one must test for continuity (which has a precise mathematical definition, see Figure S12). However, if a delay map from a putative causee to a putative causer is not continuous, we need to decide whether this is because of noise and discrete sampling, or instead because the putative causal relationship is not there (Figure 4, compare D and F).

Several methods have been used to detect SSR causal signals by detecting approximate continuity [15] or related properties [5, 45, 14]. The most popular is convergent cross mapping (CCM), which has been applied to nonlinear [5] or linear deterministic systems [35]. CCM is based on a statistic called “cross map skill”. Cross map skill quantifies how well a causer can be predicted from delay vectors of its causee (Figure 6A), conceptually similar to checking for gradual transitions when shading the causee delay space by causer values (Figure 4). Four criteria have been proposed to infer causality [5, 11, 12] (Figure 6B): First, the cross map skill must be positive. Second, the cross map skill must be significant according to some surrogate data test. Third, the cross map skill must increase with an increasing amount of training data. Lastly, cross map skill must be greater when predicting past values of causer than when predicting future values of causer (the previously mentioned prediction lag test [11, 12], but see Appendix 1.15). In practice, many if not most CCM analyses use only a subset of these four criteria [5, 49, 50, 51]. Other approaches to detect various aspects of continuous delay maps have also been proposed [45, 15, 14, 76]. We do not know of a systematic comparison of these alternatives.

## Simulation examples: External drivers and noise jointly influence causal inference performance

In this section we examine how environmental drivers, process noise, and measurement noise can influence the performance of Granger causality and CCM, using computer simulations. We constructed a toy ecological system with a known causal structure, obtained its dynamics (with noise) through simulations, and applied a linear Granger causality test (using the MVGC package [27]) and CCM (using the R language package rEDM) to test how well we can infer causal relationships.

We simulated a two-species community in which one species (*S*_1_) causally influences the other species (*S*_2_) but *S*_2_ has no influence on *S*_1_ (Figure 7B). Additionally, *S*_1_ is causally influenced by an unobserved periodic external driver and *S*_2_ either is (Figure 7D) or is not (Figure 7E) causally influenced by its own (also unobserved) periodic external driver. We added process noise to model the stochastic nature of natural ecosystems and added measurement noise to model measurement uncertainty. Process noise propagates to future time steps and can result from, for instance, stochastic migration and death (Figure 7A). In contrast, measurement noise does not propagate over time, and includes instrument noise as well as ecological processes that occur during sampling. Unlike in linear Granger causality, there is no default test procedure for CCM causality criteria [12, 35]. We therefore tested for CCM criteria using two different procedures (Figure 7 legend and Supplementary Methods).

**Figure 7:**
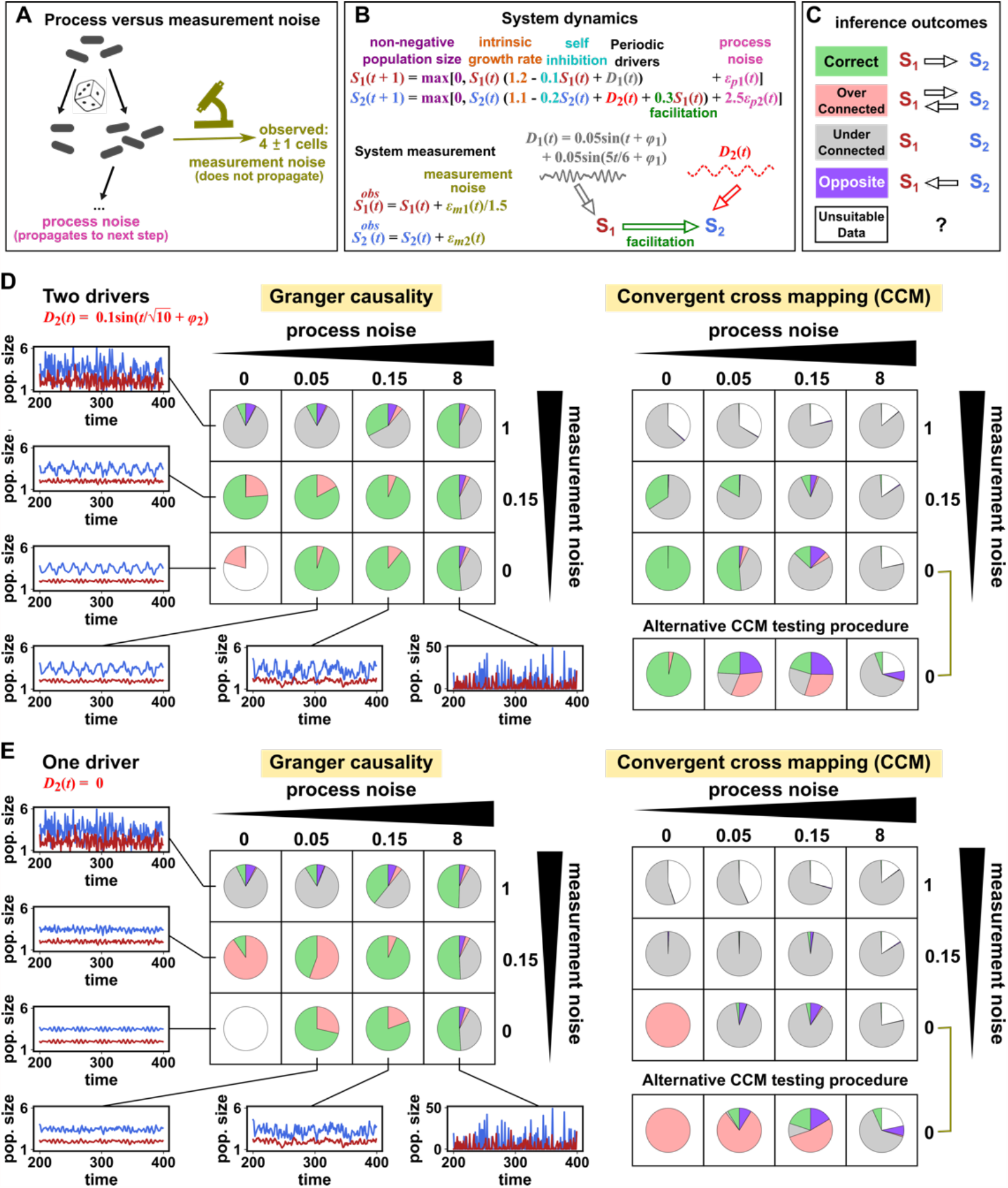
Performance of Granger causality and convergent cross mapping in a toy model with noise. (**A**) The effect of process noise but not measurement noise propagates to samples taken at subsequent time points. (**B**) We simulated a two-species community. The process noise terms *ϵ*_*p*1_(*t*) and *ϵ*_*p*2_(*t*), as well as the measurement noise terms *ϵ*_*m*1_(*t*) and *ϵ*_*m*2_(*t*), are IID normal random variables with a mean of zero and a standard deviation whose value we vary. (**C**) Five possible outcomes of causal inference. (**D, E**) Community dynamics and causal inference outcomes. We varied the level (i.e. standard deviation) of process noise and measurement noise. Each pie chart shows the distribution of inference outcomes from 1000 independent replicates. Granger causality: In the deterministic setting (lower left corner), the MVGC tool correctly rejects the data as inappropriate. With modest (but not high) process noise and no measurement noise, Granger causality frequently infers the correct causal structure. High measurement noise prevents the detection of true causality, as expected. With low process noise (i.e. 0.05), adding an intermediate level of measurement noise can increase the false positive rate. CCM: The performance of CCM is sensitive to test procedure (olive brackets) and ecological details (one versus two drivers), and quickly deteriorates with measurement or process noise. When *S*_2_ does not have its own external driver (**E**), CCM infers false causality in the deterministic case due to the (visually apparent) synchrony. In both the main and alternative CCM procedures, criterion 1 (positive *ρ*) was checked directly and random phase surrogate data were used to test criterion 2 (significance of *ρ*). Criterion 4 (prediction lag test) was not used, because the test is difficult to interpret for periodic dynamics where cross map skill can oscillate as a function of prediction lag length (Figure S16). The two procedures differ only in how they test criterion 3 (*ρ* increases with more training data): the main procedure uses bootstrap testing following [12] while the alternative procedure uses a Kendall’s *τ* as suggested by [53].

Granger causality and CCM can perform well when their respective requirements are met, but both are fairly sensitive to the levels of process and measurement noise (Figure 7D and E, correct inferences colored as green in pie charts). In reality, where a system lies in the spectrum of process versus measurement noise is often unknown, and we are not aware of any test that reliably distinguishes between process noise and measurement noise. Both Granger causality and CCM are also sensitive to details of the ecosystem (whether or not *S*_2_ has its own external driver; compare Figure 7D with E). CCM is additionally sensitive to test procedure details (Figure 7D and E, olive brackets).

## Summary: Model-free causality tests are not assumption-free

Returning to the three questions we initially posed, we can summarize answers in Fig 8. Although the techniques explored in this article are model-free in the sense that they do not depend on prior mechanistic knowledge, they are by no means free from assumptions. Such assumptions are not necessarily evils. However, the danger in replacing knowledge-based modeling with model-free inference is that we can replace explicitly stated assumptions with unstated and unscrutinized assumptions. Too frequently, both methodological and applied works fall into this trap. Nevertheless, when assumptions are clearly articulated and shown to be reasonable, model-free causal inference techniques have the potential to jump-start the discovery process where little mechanistic information is known.

**Figure 8:**
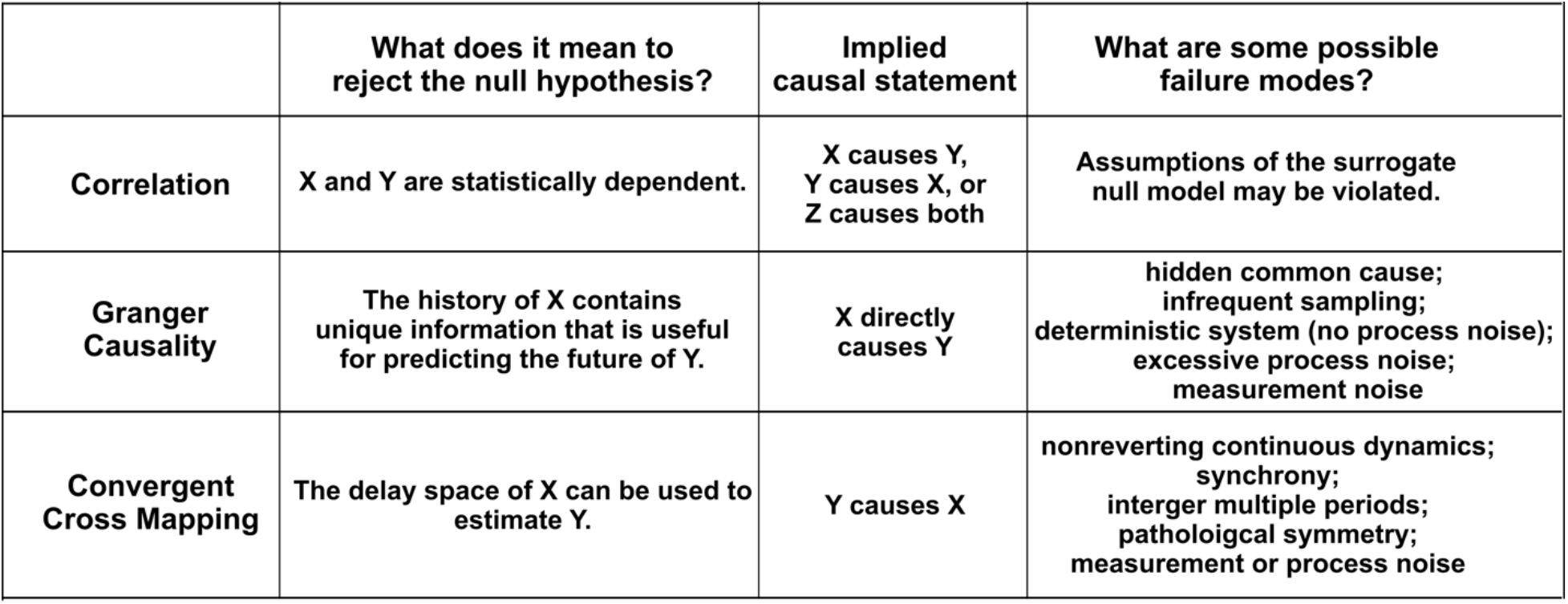
A comparison of three statistical causal inference approaches.

## Acknowledgments

Ideally, writers of this essay should be Renaissance scholars with mastery of divergent fields. However, human knowledge has expanded since the Renaissance and we are not modern da Vincis. We therefore sought feedback and advice from domain experts: Bree Cummins (Montana State University) and Tim Sauer (George Mason University) discussed topology with us; Kun Zhang (Carnegie Mellon University) held extensive discussions on causal inference with us. Kun Zhang (Carnegie Mellon University), Sean Gibbons (Institute for Systems Biology), Nathan Kutz (University of Washington), Peng Ding (UC Berkeley), Chris Barnes (UCL), Bianca De Stavola (UCL), and Ricardo Silva (UCL) critiqued our manuscript. We thank David Fredricks and Sujatha Srinivasan (Fred Hutch) for discussions that inspired this effort, and members of the Shou group for helpful comments.

