## Supplementary materials for "Data-driven causal analysis of observational time series in ecology"

#### 1 Appendices

##### 1.1 Dependence between random variables and between vectors of random variables

The random variable is a concept important both for the field of statistics as a whole, and for statistical methods that concern causality. A random variable is a variable whose values or experimental measurements depend on outcomes of a random phenomenon and follow a particular probability distribution. Reichenbach's common cause principle states that if $X$ and $Y$ are random variables and have a statistical dependence (such as a nonzero covariance), then one or more of three statements is true: $X$ perturbation causes $Y$, $Y$ perturbation causes $X$, or a third variable $Z$ perturbation causes both $X$ and $Y$. The common cause principle cannot be proven from the axioms of probability; rather, the principle is itself a fundamental assumption that supports much of the modern statistical theory of causality ([8] Section 1.4.2).

As an example, consider the sizes of cells in a microbial population. We can use random variables $X_1$ and $X_2$ to record the volume and the length of a cell, and then repeat measurements for many cells (trials) to get data (realizations). If larger cells tend to be longer, then the volume and the length random variables covary and are thus dependent. A mathematical definition of dependence (and its opposite, independence) is presented in Figure S1B. Other equivalent definitions exist (see Appendix 1.3).

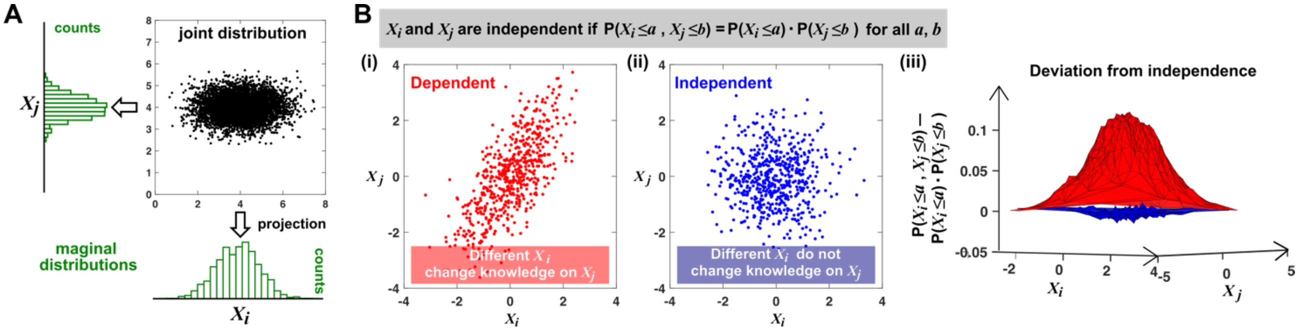


Figure S1: Joint distribution, marginal distributions, and dependence of two random variables. (**A**) A scatterplot of data associated with random variables $X_i$ and $X_j$ represents a "joint distribution" (black). Histograms for data associated with $X_i$ and for data associated with $X_j$ represent "marginal distributions" (green). Strictly speaking, joint and marginal distributions must be normalized so that probabilities (here represented as "counts") sum to 1. Graphically, marginal distributions are projections of the joint distribution on the axes. Two random variables are identically distributed if their marginal distributions are identical. (**B**) Independence between two random variables. Gray box: the mathematical definition of independence, where "$P$" means probability. Two random variables are dependent if and only if they are not independent. Visually, if two random variables are independent, then different values of one random variable will not change our knowledge about another random variable. In (**i**), $X_j$ increased as $X_i$ increased (different $X_i$ led to different knowledge on $X_j$), and thus, $X_i$ and $X_j$ are not independent (i.e. they are dependent). In (**ii**), $X_i$ and $X_j$ were repeatedly drawn from two normal distributions. Thus, the two random variables are independent. One might argue that when $X_i$ values become extreme, $X_j$ values tend to land in the middle. However, this is a visual artifact caused by fewer data points at the more extreme $X_i$ values. If we had plotted histograms of $X_j$ at various $X_i$ values, we would see that $X_j$ is always normally distributed with the same mean and variance. (**iii**) Indeed, when we plotted the difference between the observed probability $P(X_i \le a, X_j \le b)$ and the probability expected from $X_i$ and $X_j$ being independent $P(X_i \le a) \cdot P(X_j \le b)$, (ii) showed a near-zero difference (blue), while (i) showed deviation from zero (red). This is consistent with $X_i$ and $X_j$ being independent in (ii) but not in (i).

Note that when measuring a random variable, sampling is with replacement, or can be thought of as from an infinite population. For example during dice rolling, if the first trial registers 1, then the second trial can register 1 as well. Otherwise, if sampling was done *without* replacement, then the second trial would not register 1, which means that the outcome of the second trial would depend on the outcome of the first trial. This would violate the requirement that realizations of a random variable should be independent. As another example, imagine that we raised 20 mice, and sacrificed 4 mice per time point for 5 time points. Although at each time point sampling was done without replacement, the original 20 mice can be thought of as being sampled from an infinite population of possible mice. Thus any pair of mice in this experiment can be considered independent.

Dependence can be readily generalized as a property between two vectors of random variables. (Note that a time series can be viewed as a vector of random variables; Figure S7iii.) For example, suppose that we measure two variables $X$ and $Y$ over two days. Our (very short) time series are then $[X_1, X_2]$ and $[Y_1, Y_2]$ where the subscript index denotes the day of measurement. Similar to Figure S1B, we would say that our two time series are independent if

$$P(X_1 \leq x_1, X_2 \leq x_2, Y_1 \leq y_1, Y_2 \leq y_2) = P(X_1 \leq x_1, X_2 \leq x_2)P(Y_1 \leq y_1, Y_2 \leq y_2)$$

for all choices of $x_1, x_2, y_1, y_2$.

#### 1.2 When are two random variables independent and identically distributed (IID)?

Two random variables are IID if they have the same probability distribution and are independent. Figure S1 above illustrates how to check whether two random variables have identical distributions and are independent. In Figure S2 we give examples of pairs of random variables that are (or are not) identically distributed, and that are (or are not) independent. Note that two dependent random variables can be linearly correlated (Figure S2 3rd column), or not (Figure S2 4th column). A common quantitative measurement of linear correlation known as the Pearson correlation coefficient is nicely illustrated at https://en.wikipedia.org/wiki/Pearson_corre

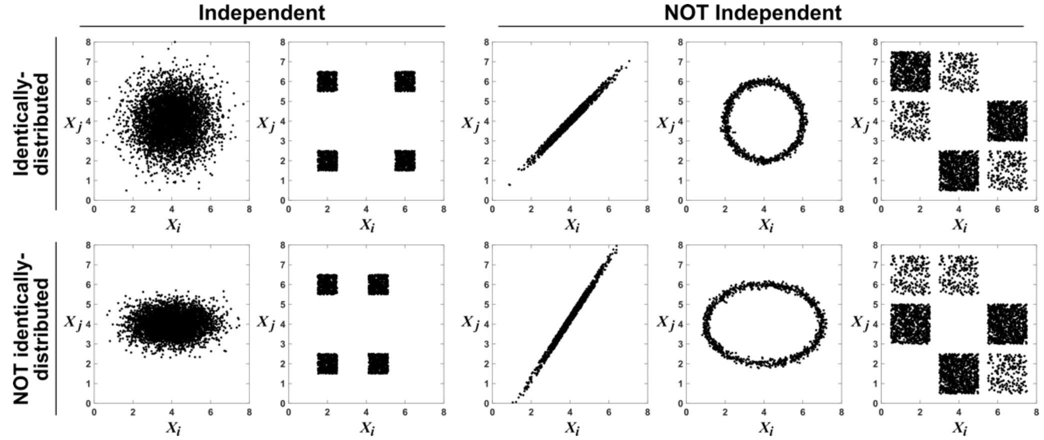


Figure S2: **Examples of random variables that are identically distributed or not identically distributed, and independent or not independent.** In the top row, $X_i$ and $X_j$ are identically distributed (projections of the scatter plot on both axes would have the same shape). Note that in the top row of the rightmost column, the scatter plot is not symmetric along the diagonal line, yet projections on both axes yield identical marginal distributions: three segments of equal densities. Thus, the two random variables are identically distributed. In the bottom row, $X_i$ and $X_j$ are not identically distributed. In the leftmost two columns, the two random variables are independent (for more details about independence, see Figure S1B). In the last three columns, the two random variables are dependent: different $X_i$ values alter our knowledge of $X_j$.

A sample of values drawn from a mixed population can still constitute an IID dataset, as long as sample members are chosen randomly and independently. For example, consider a population of replicate mice where we observe that fecal biomass measurements of Microbe 1 and Microbe 2 are correlated, and we want to know whether we can expect a causal explanation. We can attempt to satisfy the requirement of independence by ensuring that none of the mice in this study interacts with any of the other mice in this study (e.g. sampling from different cages). We can attempt to satisfy the requirement of identical distributions by imposing similar environmental and genetic conditions on each mouse so that any differences in the biology of our replicate mice are random. Moreover, suppose that we measure the load of Microbe 1 in each of several mice where female mice tend to have a higher load than male mice. Our measurements can still be IID if we measure Microbe 1 from both male and female mice, as long as we randomly choose among male and female mice instead of preassigning a male to female ratio (Figure S3). In typical situations, we cannot guarantee IID with mathematical certainty (e.g. we cannot guarantee that no two human subjects interact), so the researcher must decide whether the IID assumption is adequate.

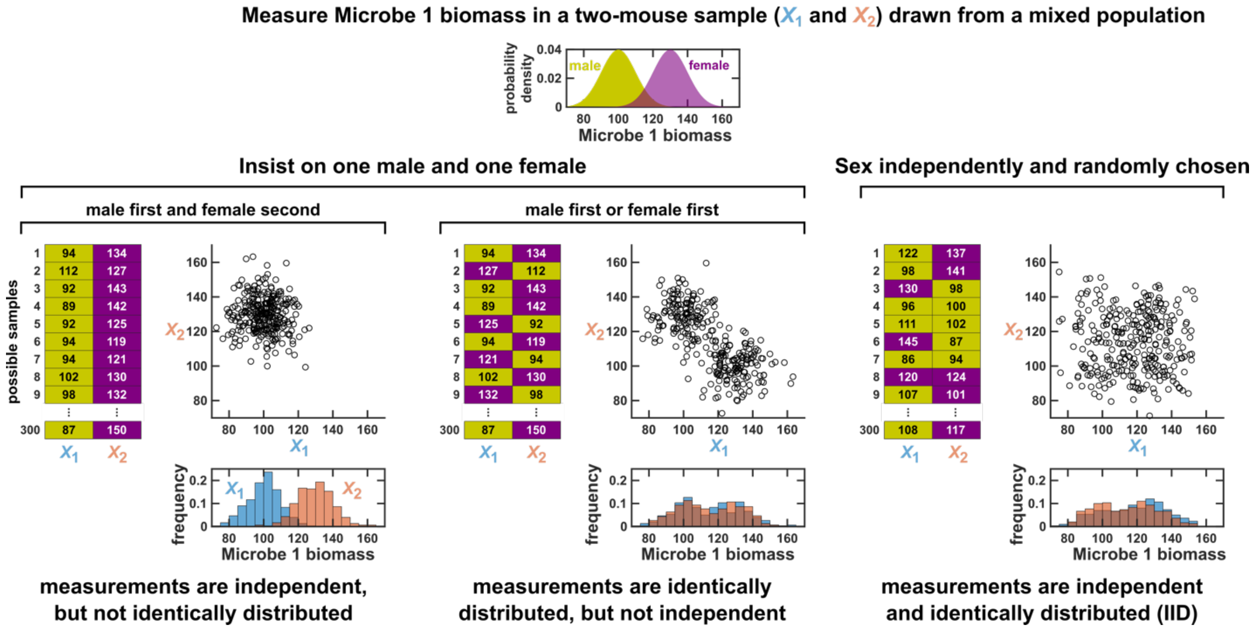


Figure S3: **Measurements taken from a mixed population may still be IID, as long as sampling is independent and random.** Consider a study in which we measure Microbe 1 biomass from a mixed population of low-load male mice and high-load female mice. For simplicity, suppose that each sample comprises measurements of two mice. To see whether a sample could be an IID dataset, we imagine drawing many possible versions of that sample, and ask whether our first measurement $X_1$ and second measurement $X_2$ are identically distributed and independent across possible samples. We could draw this sample in 3 different ways (3 sets of charts). On the left, we take our first measurement from a male and second measurement from a female. In this case, our two measurements are independent, but not identically distributed, so our sample is not an IID dataset. In the middle, we choose one male and one female per sample, but choose the first measurement randomly from a male or female. Now, our measurements are identically distributed (save for sampling error) but not independent (so also not IID). On the right, the sex of each measurement is randomly and independently chosen, and thus a sample could have two measurements from the same sex. In this case our sample is an IID dataset.

#### 1.3 Conditional independence

In Appendix 1.1, we discussed independence between two random variables. Here, we will restate the concept of independence in equivalent forms that will allow us to more easily transition to the concept of conditional independence.

It is intuitive that two variables are independent if knowledge of one variable tells us nothing about the other. The statistical notion of independence captures this intuition: Random variables $X$ and $Y$ are independent if the conditional distribution of $X$ given $Y$ is always equal to the marginal distribution of $X$. For discrete random variables, this condition can be written $P(X = x|Y = y) = P(X = x)$, or equivalently $P(X = x, Y = y) = P(X = x)P(Y = y)$. For continuous random variables, independence can be written in terms of probability density functions as $f_X(x|y) = f_X(x)$ or equivalently, $f_{X,Y}(x, y) = f_X(x)f_Y(y)$ where $f_X(x|y)$ is the conditional density of $X$ given $Y$, $f_{X,Y}(x, y)$ is the joint density of $X$ and $Y$, and $f_X(x)$ and $f_Y(y)$ are the marginal densities of $X$ and $Y$, respectively (Appendix 1.1).

The statement "$X$ and $Y$ are conditionally independent given $Z$" intuitively means that $X$ and $Y$ are independent when we only analyze outcomes where $Z$ has a certain value. For discrete random variables, this condition is written $P(X = x|Y = y, Z = z) = P(X = x|Z = z)$, or equivalently, $P(X = x, Y = y|Z = z) = P(X = x|Z = z)P(Y = y|Z = z)$. For continuous random variables, we have a similar formulation except that probability $P$ is replaced by probability density $f$ (e.g. $f_{X,Y}(x, y|z) = f_X(x|z)f_Y(y|z)$ for all $x, y, z$). If $X$ and $Y$ are not conditionally independent given $Z$, then $X$ and $Y$ are conditionally dependent given $Z$.

#### 1.4 The significance of a correlation between time series can be relatively easily tested when multiple trials exist

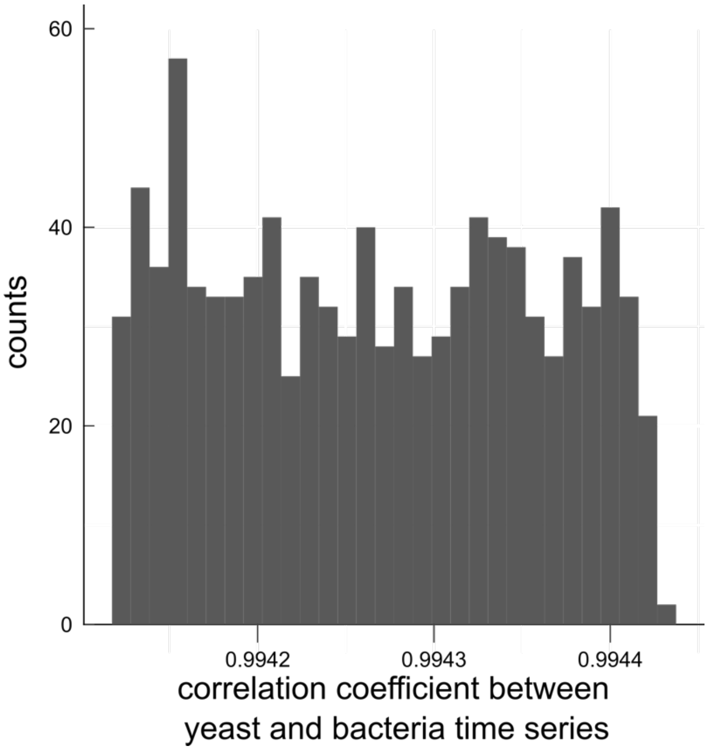


Figure S4: Independent time series may produce a high Pearson correlation coefficient, but this correlation will likely be unimpressive when compared to the appropriate null model. Suppose that yeast and bacteria grow exponentially and independently in a test tube (no resource competition or other forms of interactions). In this particular example, yeast grows with a rate of 1 while bacteria grows with a rate drawn from a uniform distribution between 1.3 and 1.31. Within one of these trials, the Pearson correlation coefficient between yeast and bacteria time series (21 time points from 0 to 20) is 0.9941. Although this coefficient is nearly 1, it is not impressive when compared to the histogram of Pearson correlation coefficients obtained when correlating the same yeast series with bacteria time series from different tubes (shown above).

#### 1.5 Inferring causal relations and their associated directed acyclic graphs (DAGs)

It is common in causal analysis to construct a graph to represent causal relationships (a causal graph). The graph consists of vertices (random variables) and directed edges pointing from a direct cause (or "Markovian parent", Figure 1B) to its causee (or child). A directed acyclic graph (DAG) is a directed graph with no directed cycles (i.e. devoid of any directed paths from a variable to itself; Figure S5ii). Acyclicity is commonly required for nice theoretical properties and ease of analysis [19]. Additionally, when data are temporal, a particular node in the graph commonly refers to a particular variable measured at a particular time [2]. If we follow this convention and note that causation cannot flow backward in time, and if we additionally exclude instantaneous causation, then our causal graph will be acyclic, even for systems with feedback (Figure S7).

DAGs are useful visual tools, but for many purposes we need to be more mathematically precise about what we mean when we draw an edge from one variable to another. Thus, often one interprets a causal DAG as corresponding to a set of equations with the following two conditions: First, each variable can be written as a function of (only) the variable's direct causers and a random process noise term unique to the variable. Models that satisfy this condition are called structural equation models (SEMs) [17]. Second, all process noise terms are (jointly) independent of one another. SEMs that satisfy this second condition are called Markovian and have a useful property called the "causal Markov assumption" [8]. (Some authors [2], but not all [17], require that all SEMs be Markovian by definition.) The causal Markov assumption, along with the related "causal faithfulness assumption" are key tools that allow one to infer aspects of causal structure directly from data.

The causal Markov assumption can be stated in several ways that are equivalent for SEMs with independent process noise terms [2, 17]. One version states that if there is no path from $X$ to $Y$ (we cannot go from $X$ to $Y$ by following a sequence of arrows in the forward direction), then $X$ and $Y$ are conditionally independent given $X$'s Markovian parents. That is, when $X$'s direct causes are fixed, then dependence between $X$ and $Y$ implies a path from $X$ to $Y$. Such a path from $X$ to $Y$ would in turn imply that $X$ causes $Y$ in most typical cases (e.g. no cancellation of the kind in Figure 1C). Technically, $Y$ in this definition can be either a variable or a set of variables. An imprecise shorthand for this idea is "dependence implies causality". Note that if $X$ does not have any Markovian parents, then the statement "$X$ and $Y$ are conditionally independent given $X$'s Markovian parents" reduces to "$X$ and $Y$ are independent". When data have selection bias, then dependence can be illusionary, and thus any inferred causal relationship can also be illusionary (Figure S6).

The causal faithfulness assumption is, like the Markov assumption, very useful in causal inference and often quite reasonable. However, faithfulness is more difficult to state without first introducing technical notation ("$d$-separation" [2, 17]) or terminology ("entailing" [22]). We attempt to give the gist of the idea here and direct readers to other sources [22, 2] for more precise definitions. The causal faithfulness assumption is a kind of converse to the causal Markov assumption. Recall that the causal Markov assumption requires certain conditional (or unconditional) independence relationships based on the causal graph structure. Let us call any other independence relationships (i.e. those not required directly or indirectly by the causal Markov assumption) "extra" independence relationships. A graph is causally faithful to the probability distribution of its random variables if no "extra" independence relationships exist [17]. An imprecise shorthand for this idea is "independence implies lack of causality". The faithfulness assumption can be violated when two effects precisely cancel each other (e.g. Figure 1 C). Another instance where observed independence does not imply lack of causality can occur when quantities are related by saturable functions. For example, suppose that a microbe requires a certain nutrient to grow, and that the nutrient is available in vast excess of what is needed for maximal growth. Then, small fluctuations in nutrient availability could be independent of microbe growth rate, even though nutrient availability causally influences microbe growth rate (because removing the nutrient entirely would stop growth).

Existing causal discovery methods for observational nontemporal data are diverse. Such methods can differ greatly in the assumptions they make (e.g. whether there are hidden variables or "unknown shift interventions") [23]), the lines of reasoning they employ, and the resolution of causal detail they provide (e.g. a unique causal graph versus a set of several plausible graphs) [23]. We will briefly introduce two classes of causal methods: (1) constraint-based search and (2) structural equation models (SEMs) with assumptions about the functional forms of equations [19]. However, these two classes, while illustrative of different modes of causal discovery, are far from an exhaustive list [23].

In constraint-based search, an algorithm (such as PC, FCI, or Boolean Satisfiability Solver) uses independence and dependence relationships (and their conditional counterparts) to narrow down the scope of causal graphs without exhaustively checking all possibilities (which can be enormous in number even for a handful of variables). However, there are often multiple graphs consistent with the data (e.g. the second to last row of Figure S5 ii, see legend).

In the functional form-based (or SEM-based) approach to causal discovery, one begins by assuming that the causal relationship follows a certain functional form such as the linear non-Gaussian acyclic model, nonlinear additive noise model, or the post-nonlinear model. Then, one tests a particular causal relationship by regressing a potential causee against its potential direct causers, and assessing whether the residual is independent from all potential direct causers. In many cases, non-Gaussian noise enables a higher resolution of causal inference.

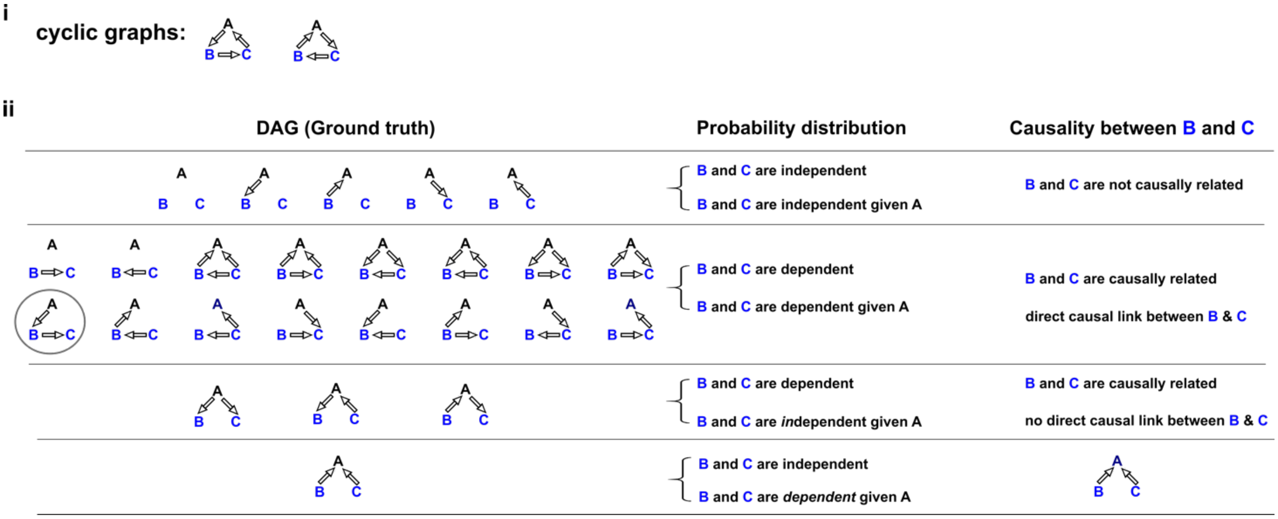


Figure S5: Probability distributions alone can specify causal structure to varying degrees of resolution. Consider a system of 3 and only 3 random variables $A, B$, and $C$. Between each of the three pairs of variables, there are in principle three possible relationships: no edge, an edge pointing from one to the other, and an edge in the opposite direction. Since we have three pairs of variables with three types of relationships, there are $3^3 = 27$ possible graphs. (**i**) Two of these are cyclic, and thus not DAGs, while the rest (ii) are DAGs. (**ii**) Left column: Ground truth causal DAGs. Center column: When a DAG is associated with a Markovian and causally faithful SEM, we can infer some aspects of causal structure from the probability distributions alone. We demonstrate this by using the probability distributions (center column) of $B$ and $C$ (blue) to infer causal links between $B$ and $C$ (right column). Top row: $B$ and $C$ are independent. $B$ and $C$ are not causally related. Second row: $B$ and $C$ are dependent, implying that they are causally related. Furthermore, $B$ and $C$ are conditionally dependent given $A$. For example, in the circled graph, for a given value of $A$, variation in $B$ will affect $C$, leading to dependence between $B$ and $C$. Indeed, there is a direct link between $B$ and $C$. Third row: $B$ and $C$ are dependent, but become independent given $A$. In this case, there is no direct link between $B$ and $C$, but they are causally related. Note that all three graphs are consistent with the following observations: $B$ and $C$ are conditionally independent given $A$; $A$ and $C$ are conditionally dependent given $B$; $A$ and $B$ are conditionally dependent given $C$. Thus, we cannot uniquely identify the causal structure from probability distributions alone. Last row: $B$ and $C$ are independent, but become dependent upon conditioning on $A$. There is a unique causal graph corresponding to this scenario; see Figure S6 for an example (math and writing scores are independent, but become dependent when conditioned on college admission).

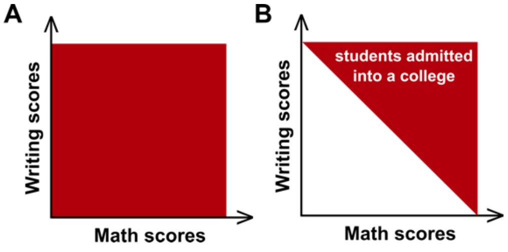


Figure S6: Selection bias creates false dependence. (A) Math and writing scores in the general student population are independent of each other. (B) A college admits students with a total score of higher than a fixed number. In this selected dataset of admitted students, math and writing scores are no longer independent.

#### 1.6 Time series of systems with feedback loops can be analyzed by methods designed for directed acyclic graphs

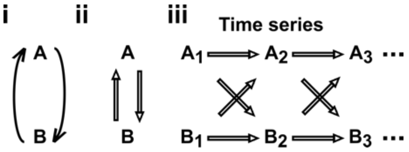


Figure S7: Causal inference approaches designed for directed acyclic graphs (DAGs) can be applied to time series from systems with feedback. (**i**) Consider a mutualistic system where $A$ and $B$ represent the population sizes of two species that mutually facilitate each other's growth. (**ii**) For data without temporal information, the causal graph is cyclic and thus not a DAG. (**iii**) For time series data where $A_1$, $A_2$, ... represent the population size of $A$ at times $1, 2, ...$, the causal graph is no longer cyclic since $A_1$ causes $B_2$ and $B_1$ causes $A_2$ etc. Note that $A_1$ causes $A_2$ (and similarly $B_1$ causes $B_2$). This framework [2] has helped the authors classify mutations in communities with feedback [20, 21].

#### 1.7 Intuition for random phase test

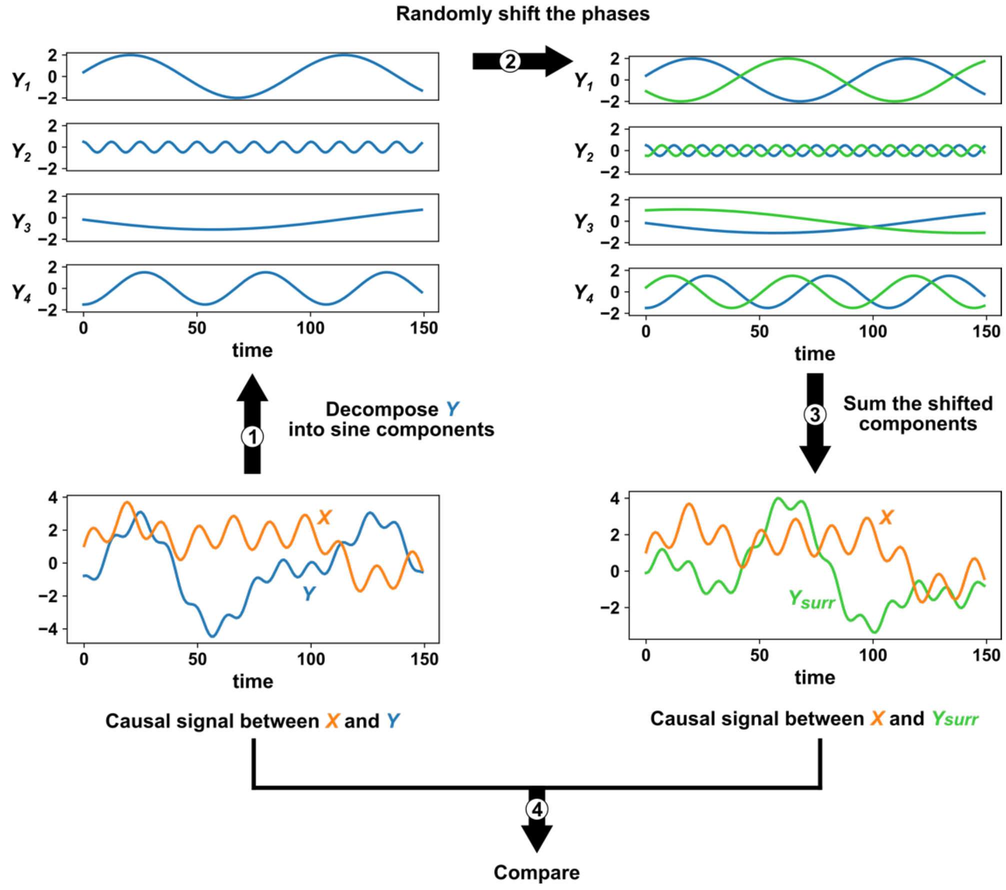


Figure S8: Intuition for random phase surrogate data methods. Given two time series $X$ and $Y$ (lower left), we can compute a correlation statistic to detect dependence. Surrogate data methods test the significance of this correlation by comparing it to correlations generated when one of the time series ($Y$) is replaced by a similar time series ($Y_{surr}$) which is assumed to be independent of $X$. Random phase surrogate data methods generate $Y_{surr}$ by representing $Y$ as a sum of sine waves (upper left), randomly shifting the phases of the component sine waves (upper right green), and summing up the shifted sine waves (lower right green). A $p$-value is calculated as the fraction of $(X, Y_{surr})$ time series that produce a causal signal that is at least as strong as the causal signal from the original $(X, Y)$ time series.

#### 1.8 Covariance-stationarity

A stochastic process $X_t$ is covariance-stationary (or wide-sense stationary) if:

1. $\mathbb{E}[\mathrm{X_t}]$ (the ensemble mean) does not depend on $t$
2. $\mathrm{Var}[X_t]$ is finite and does not depend on $t$
3. For all choices of $h$, $\mathrm{Cov}(X_t, X_{t+h})$ does not depend on $t$

As an example similar to Figure 2C, consider a population whose dynamics are governed by death and stochastic migration:

$$X_t = (1 - a)X_{t-1} + c + \epsilon_t \tag{3}$$

Here, $X_t$ is the population size at time $t$, $a$ is the probability of death during the time interval of 1, $c$ is the average number of individuals migrating into the population during the time interval of 1, and $\epsilon_t$ is a random variable with a mean of zero which represents temporal fluctuations in the number of migrants. Suppose that we observed the dynamics of 10 populations governed by Eq. 3 such that the populations all have the same parameters, but are independent (FigS9A). Then, at each time point $t$, we will have some distribution of values of $X_t$. In fact, if we have not just 10, but 1,200 replicates, we can see that the distribution of values of $X_t$ does not appear to depend on time (Figure S9B, top). Furthermore, the covariance between $X_t$ and $X_{t+1}$ does not appear to depend on time either (Figure S9B, bottom).

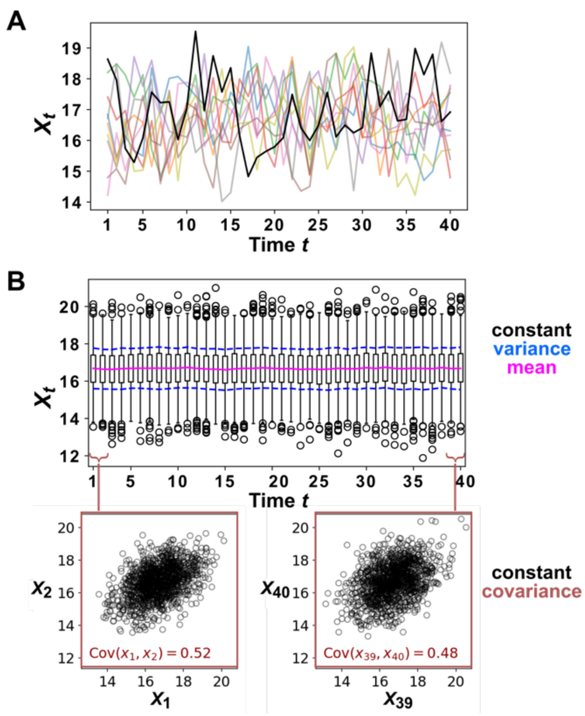


Figure S9: Example of a stationary process. (**A**) Ten replicate runs of the stochastic process described in Eq. 3 with parameter choices $a = 0.6$, $c = 10$, and $\epsilon$ is a normal random variable with mean of zero and standard deviation of 1. (**B**) The distribution $X_t$ values shown for $1,200$ replicates runs of the same stochastic process as in (A). The mean of $X_t$ is given as a solid red line and the mean $\pm$ standard deviation of $X_t$ is shown as a dashed blue line. Bottom: $X_t$ is plotted against $X_{t+1}$ for two values of $t$.

Although it is common to talk about a time series being stationary or nonstationary, this is technically a slight abuse of language. Just as the mean and variance are properties of a random variable (and not of any single data point obtained from that random variable), stationarity is a property of a stochastic process (and not of any single time series produced by that process). This fact is illustrated by comparing the middle and bottom rows of Figure S10. If we examine any one time series from the middle or bottom rows (e.g. the black curves in each), we see that they have essentially the same dynamics (i.e. they are sine waves with the same frequency). However, the process shown in the middle row is covariance-stationary (as shown below), whereas the process shown in the bottom row is not since its mean changes over time.

To see that the middle row of Figure S10 shows a covariance-stationary process, we can show that the mean, variance, and covariance of the process are constant:

$$\mathbb{E}[X_2(t)] = \int_0^{2\pi} \cos\left(\frac{t}{3} + \theta_2\right) d\theta_2 = 0$$

$$\mathrm{Var}[X_2(t)] = \int_0^{2\pi} \left(\cos\left(\frac{t}{3} + \theta_2\right)\right)^2 d\theta_2 = \pi$$

$$\mathrm{Cov}(X_2(t), X_2(t+h)) = \int_0^{2\pi} \cos\left(\frac{t}{3} + \theta_2\right) \cos\left(\frac{t+h}{3} + \theta_2\right) d\theta_2 = \pi \cos\left(\frac{h}{3}\right)$$

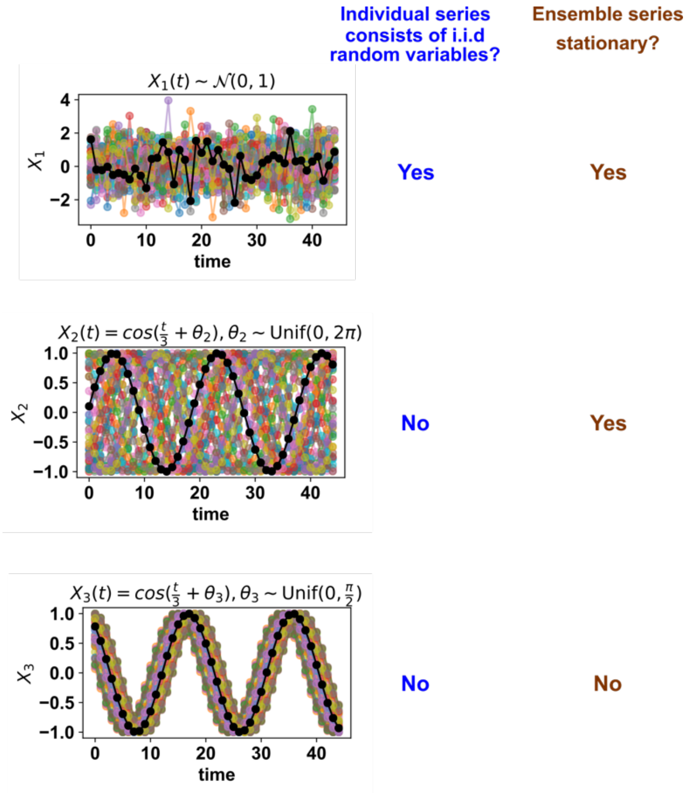


Figure S10: Whether a stochastic process is stationary depends on its ensemble of time series. The top panel shows IID standard normal noise. The middle and bottom panel both show sinusoidal curves. Although an individual time series from the middle panel looks similar to that from the bottom panel, only the middle panel shows a covariance-stationary process.

#### 1.9 Deterministic processes with many variables may appear stochastic

A deterministic time series from a system with many variables can be approximated as stochastic. This is illustrated below in Figure S11. When we track the trajectory of a particle in a box with 99 other particles (Figure S11 bottom row), the observed trajectory appears random, even though the governing equations of motion are deterministic. In particular, the motion of our particle over each time step can be approximated as having a random component. This sets the apparent randomness of many-variable dynamics apart from that of a different phenomenon called chaos. In chaotic dynamics, each time step needs not be random, but small changes in initial conditions lead to large changes at later times.

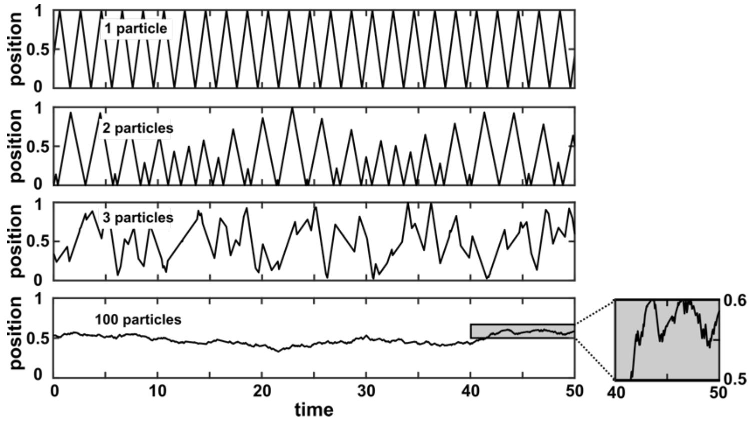


Figure S11: A many-variable deterministic system can be approximated as a stochastic system. The position of a particle in a system of particles bouncing in a 1-dimensional box is plotted over time. In each simulation, particles with radius 0 bounced in a box with walls of infinite mass placed at positions 0 and 1. All particles have mass 1 and are initialized at a random position between 0 and 1 according to a uniform distribution. Initial velocities are chosen in the following way: The initial velocity of each particle in a box is first randomly chosen from between −1 and 1 according to a uniform distribution. Then, all initial velocities in a given box are multiplied by the same constant to ensure that the total kinetic energy of each box is 0.5. Kinetic energy is conserved throughout the simulation. The simulation then follows the particles as they experience momentum-conserving collisions with one another and with the walls.

#### 1.10 Infrequent sampling

In this section, we consider the systems shown in Figure 3Biv and show how one can derive the infrequent sampling case from the frequent sampling case. In Figure 3Biv we show the system

$$X_t = 0.4X_{t-1} + 0.6Y_{t-1} + \epsilon_{X,t} \tag{4}$$
$$Y_t = 0.5Y_{t-1} + \epsilon_{Y,t}$$

where $\epsilon_{X,t}$ and $\epsilon_{Y,t}$ are independent normal random variables with 0 mean and standard deviation of 1. We state that if one samples this system infrequently (every 10 time steps), one arrives at the following system

$$X_t \approx 0.0001X_{t-10} + 0.005Y_{t-10} + 1.431\beta_{X,t} \tag{5}$$
$$Y_t \approx 0.001Y_{t-10} + 1.155\beta_{Y,t}$$
$$\text{Cov}(\beta_{X,t}, \beta_{Y,t}) \approx 0.303$$

where $\beta_{X,t}$ and $\beta_{Y,t}$ are normal random variables with 0 mean and standard deviation of 1. This demonstrates that decreasing the sampling frequency can reduce the ratio of signal (effect of causer) to noise (effect of process noise). We now show how one can derive Eq. 5 from Eq. 4.

We begin by rewriting Eq. 4 in matrix form and putting $t+1$ rather than $t$ on the left hand side:

$$\begin{pmatrix} X_{t+1} \\ Y_{t+1} \end{pmatrix} = \begin{pmatrix} 0.4 & 0.6 \\ 0 & 0.5 \end{pmatrix} \begin{pmatrix} X_t \\ Y_t \end{pmatrix} + \begin{pmatrix} \epsilon_{X,t+1} \\ \epsilon_{Y,t+1} \end{pmatrix}$$

In general, this looks like:

$$\boldsymbol{U}_{t+1} = \boldsymbol{A}\boldsymbol{U}_t + \boldsymbol{\epsilon}_{t+1} \tag{6}$$

Here, for a system of $m$ state variables, $\boldsymbol{U}_t$ is a $m \times 1$ vector describing variables of the system at time $t$, $\boldsymbol{A}$ is an $m \times m$ matrix describing how the system evolves, and $\boldsymbol{\epsilon}_t$ is an $m \times 1$ vector-valued random variable that introduces process noise to the system. Eq. 6 looks one step ahead. We can look two steps ahead by plugging Eq. 6 into itself.

$$\begin{aligned} \boldsymbol{U}_{t+2} &= \boldsymbol{A}\boldsymbol{U}_{t+1} + \boldsymbol{\epsilon}_{t+2} \\ &= \boldsymbol{A}^2\boldsymbol{U}_t + \boldsymbol{A}\boldsymbol{\epsilon}_{t+1} + \boldsymbol{\epsilon}_{t+2} \end{aligned}$$

More generally, we can look $n$ steps ahead (as if we were sampling every $n$ steps rather than every step) according to

$$\boldsymbol{U}_{t+n} = \boldsymbol{A}^n\boldsymbol{U}_t + \sum_{j=1}^{n} \boldsymbol{A}^{n-j}\boldsymbol{\epsilon}_{t+j} \tag{7}$$

A proof of Eq. 7 is provided at the end of this section. Let us now assume that $\boldsymbol{\epsilon}_t$ $(t = 1, 2, ...)$ are IID multivariate normal distributions with mean vector $\boldsymbol{\mu}$ and covariance matrix $\boldsymbol{C}$. For example, if $\epsilon$ is the IID standard normal distribution, then $\boldsymbol{\mu}$ is the $m \times 1$ zero vector and $\boldsymbol{C}$ is the $m \times m$ identity matrix whose diagonal entries are 1s and whose off-diagonal entries are 0s. In general, Eq. 7 can be rewritten as:

$$\boldsymbol{U}_{t+n} = \boldsymbol{A}^n \boldsymbol{U}_t + \boldsymbol{\eta}_{n,t+n} \tag{8}$$

where $\boldsymbol{\eta}_{n,t}$ is a multivariate normal variable with mean $\boldsymbol{\mu}_n$ and covariance $\boldsymbol{C}_n$ given by:

$$\begin{aligned} \boldsymbol{\mu}_n &= \left( \sum_{j=1}^{n} \boldsymbol{A}^{n-j} \right) \boldsymbol{\mu} \\ \boldsymbol{C}_n &= \sum_{j=1}^{n} \boldsymbol{A}^{n-j} \boldsymbol{C} \left( \boldsymbol{A}^{n-j} \right)^\top \end{aligned} \tag{9}$$

where $M^\top$ is the transpose of $M$. The top line of Eq. 9 follows from the linearity of expectation. The bottom line of Eq. 9 follows from the covariance rules for linear transformation (Eq. 10) or sum (Eq. 11) of vector-valued random variables.

If we plug the relevant parameters for Eq. 4 (i.e. $A = \begin{pmatrix} 0.4 & 0.6 \\ 0 & 0.5 \end{pmatrix}$, $\boldsymbol{\mu} = \begin{pmatrix} 0 \\ 0 \end{pmatrix}$, $\boldsymbol{C} = \begin{pmatrix} 1 & 0 \\ 0 & 1 \end{pmatrix}$), and $n = 10$, then $A^{10} = \begin{pmatrix} 0.0001 & 0.0052 \\ 0 & 0.0010 \end{pmatrix}$ and $\boldsymbol{C}_n \approx \begin{pmatrix} 2.048 & 0.500 \\ 0.500 & 1.333 \end{pmatrix}$. The diagonal entries of $\boldsymbol{C}_n$ are variances and the off-diagonal entries are identical and represent the covariance between the two elements of $\boldsymbol{\eta}_{n,t}$, the process noises of $X$ and $Y$ accumulated over the $n = 10$ steps since $t - n$. Finally, we rescale the noise terms in 5 by letting

$$\begin{bmatrix} \sqrt{2.048}\beta_{X,t} \\ \sqrt{1.333}\beta_{Y,t} \end{bmatrix} = \boldsymbol{\eta}_{n,t}.$$

This rescaling gives us noise terms ($\beta$) in Eq. 5 with unit variance, putting them on the same scale as the $\epsilon$ noise terms in Eq. 4. Note that now, the covariance of $\beta_{X,t}$ and $\beta_{Y,t}$ is:

$$\begin{aligned} \mathrm{Cov}(\beta_{X,t}, \beta_{Y,t}) &\approx \mathrm{Cov}\Big(\frac{\boldsymbol{\eta}_{n,t}(1)}{\sqrt{2.048}}, \frac{\boldsymbol{\eta}_{n,t}(2)}{\sqrt{1.333}}\Big) \\ &\approx \mathrm{E}\left[ \frac{1}{\sqrt{2.048}} \left( \boldsymbol{\eta}_{n,t}(1) - \mathrm{E}\left[\boldsymbol{\eta}_{n,t}(1)\right] \right) \frac{1}{\sqrt{1.333}} \left( \boldsymbol{\eta}_{n,t}(2) - \mathrm{E}\left[\boldsymbol{\eta}_{n,t}(2)\right] \right) \right] \\ &\approx \frac{\mathrm{Cov}(\boldsymbol{\eta}_{n,t}(1), \boldsymbol{\eta}_{n,t}(2))}{\sqrt{2.048}\sqrt{1.333}} \\ &\approx \frac{0.500}{\sqrt{2.048}\sqrt{1.333}} \\ &\approx 0.303 \end{aligned}$$

and Eq. 8 becomes Eq. 5.

##### Covariance of a linear transformation of a vector-valued random variable

Suppose $\boldsymbol{X}$ is an $m \times 1$ vector of random variables whose mean vector is the $m \times 1$ vector $\boldsymbol{\mu_X}$ and whose covariance is the $m \times m$ matrix $\boldsymbol{C_X}$ . Let $\boldsymbol{Y} = \boldsymbol{AX}$ where $\boldsymbol{A}$ is an $m \times m$ matrix. Thus, $\boldsymbol{Y}$ is an $m \times 1$ vector. Denote the mean and covariance of $\boldsymbol{Y}$ by $\boldsymbol{\mu_Y}$ and $\boldsymbol{C_Y}$. Then:

$$\boldsymbol{C_Y} = \boldsymbol{A C_X A}^\top \tag{10}$$

To see this, first note that $\boldsymbol{\mu_Y} = \boldsymbol{A\mu_X}$. Then we can derive Eq. 10 directly from the definition of the covariance matrix.

$$\begin{aligned}
\boldsymbol{C_Y} &= \mathrm{E}\left[(\boldsymbol{Y} - \boldsymbol{\mu_Y})(\boldsymbol{Y} - \boldsymbol{\mu_Y})^\top\right] \\
&= \mathrm{E}\left[(\boldsymbol{AX} - \boldsymbol{A\mu_X})(\boldsymbol{AX} - \boldsymbol{A\mu_X})^\top\right] \\
&= \mathrm{E}\left[\boldsymbol{A}(\boldsymbol{X} - \boldsymbol{\mu_X})(\boldsymbol{X} - \boldsymbol{\mu_X})^\top \boldsymbol{A}^\top\right] \\
&= \boldsymbol{A}\left(\mathrm{E}\left[(\boldsymbol{X} - \boldsymbol{\mu_X})(\boldsymbol{X} - \boldsymbol{\mu_X})^\top\right]\right)\boldsymbol{A}^\top \\
&= \boldsymbol{A C_X A}^\top
\end{aligned}$$

##### Covariance of a sum of independent vector-valued random variables

Suppose that $\boldsymbol{X}$ is an $m \times 1$ random variables with covariance matrix $\boldsymbol{C_X}$ and that $\boldsymbol{Y}$ is an $m \times 1$ random variables with covariance matrix $\boldsymbol{C_Y}$. Suppose further that $\boldsymbol{X}$ and $\boldsymbol{Y}$ are independent of each other. That is, each element of $\boldsymbol{X}$ is independent of each element of $\boldsymbol{Y}$. Let $\boldsymbol{C_{X+Y}}$ be the covariance matrix of $\boldsymbol{X} + \boldsymbol{Y}$. Then it is a fact that:

$$\boldsymbol{C_{X+Y}} = \boldsymbol{C_X} + \boldsymbol{C_Y} \tag{11}$$

We can see that Eq. 11 is true by showing that it holds for each element of the matrix $\boldsymbol{C_{X+Y}}$. Specifically, we will apply the definition of covariance to the element of $\boldsymbol{C}_{\boldsymbol{X}+\boldsymbol{Y}}$ at the $i$th row and the $j$th column:

$$\begin{aligned}
(\boldsymbol{C}_{\boldsymbol{X}+\boldsymbol{Y}})_{ij} &= \mathrm{Cov}(X_i + Y_i, X_j + Y_j) \\
&= \mathrm{E}\left[\left((X_i - \mathrm{E}\left[X_i\right]) + (Y_i - \mathrm{E}\left[Y_i\right])\right)\left((X_j - \mathrm{E}\left[X_j\right]) + (Y_j - \mathrm{E}\left[Y_j\right])\right)\right] \\
&= \mathrm{E}[(X_i - \mathrm{E}\left[X_i\right])(X_j - \mathrm{E}\left[X_j\right]) + (X_i - \mathrm{E}\left[X_i\right])(Y_j - \mathrm{E}\left[Y_j\right]) + \\
&\qquad (Y_i - \mathrm{E}\left[Y_i\right])(X_j - \mathrm{E}\left[X_j\right]) + (Y_i - \mathrm{E}\left[Y_i\right])(Y_j - \mathrm{E}\left[Y_j\right])] \\
&= \mathrm{Cov}(X_i, X_j) + \mathrm{Cov}(X_i, Y_j) + \mathrm{Cov}(Y_i, X_j) + \mathrm{Cov}(Y_i, Y_j) \\
&= \mathrm{Cov}(X_i, X_j) + \mathrm{Cov}(Y_i, Y_j) \\
&= (\boldsymbol{C}_{\boldsymbol{X}})_{ij} + (\boldsymbol{C}_{\boldsymbol{Y}})_{ij}
\end{aligned}$$

where the second to last line uses the fact that $\mathrm{Cov}(Y_i, X_j) = \mathrm{Cov}(X_i, Y_j) = 0$ since $\boldsymbol{X}$ and $\boldsymbol{Y}$ are independent of each other.

#### Proof of Eq. 7

We prove Eq. 7 by induction. For the base case, let $n = 1$. Then Eq. 7 reduces to Eq. 6. For the inductive step, we assume Eq. holds for $n = k$ and wish to show that it holds for $n = k + 1$. Thus,

$$\begin{aligned}
\boldsymbol{U}_{t+k+1} &= \boldsymbol{A}(\boldsymbol{U}_{t+k}) + \boldsymbol{\epsilon}_{t+k+1} \\
&= \boldsymbol{A}\left(\boldsymbol{A}^k \boldsymbol{U}_t + \sum_{j=1}^{k} \boldsymbol{A}^{k-j} \boldsymbol{\epsilon}_{t+j}\right) + \boldsymbol{\epsilon}_{t+k+1} \\
&= \boldsymbol{A}^{k+1} \boldsymbol{U}_t + \sum_{j=1}^{k} \boldsymbol{A}^{k-j+1} \boldsymbol{\epsilon}_{t+j} + \boldsymbol{\epsilon}_{t+k+1} \\
&= \boldsymbol{A}^{k+1} \boldsymbol{U}_t + \left(\boldsymbol{A}^k \boldsymbol{\epsilon}_{t+1} + \boldsymbol{A}^{k-1} \boldsymbol{\epsilon}_{t+2} + \cdots + \boldsymbol{A} \boldsymbol{\epsilon}_{t+k}\right) + \boldsymbol{\epsilon}_{t+k+1} \\
&= \boldsymbol{A}^{k+1} \boldsymbol{U}_t + \sum_{j=1}^{k+1} \boldsymbol{A}^{k+1-j} \boldsymbol{\epsilon}_{t+j}
\end{aligned}$$

The final line shows that Eq. 7 holds for $n = k + 1$, completing the inductive step and hence the proof.

#### 1.11 Difficulty of evaluating the continuity or smoothness of a function with finite or noisy data

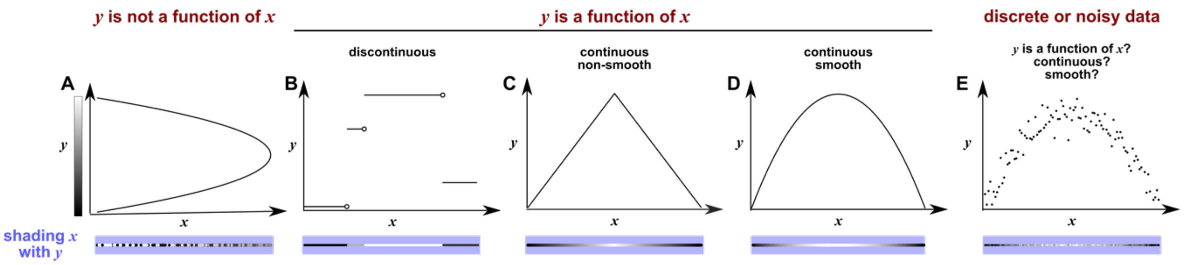


Figure S12: Continuity, smoothness, and the difficulty of evaluating the continuity or smoothness of a function with finite or noisy data. (**A**) $y$ is not a function of $x$ because a single $x$ value can correspond to more than one $y$ value. Here, when we shade $x$ with $y$ value, we randomly choose the upper or the lower $y$ value, leading to bumpy shading, similar to what we might expect to occur in the real world. (**B**) $y$ is a discontinuous function of $x$. This is because at any "breakpoint" (circle) between two adjacent segments, the limit taken from the left-hand side is unequal to the limit taken from the right-hand side. Shading $x$ with $y$ generates a "bumpy" pattern. (**C**) $y$ is a continuous and nonsmooth function of $x$. The function is nonsmooth because at the maximum point, the slope taken from the left-hand side is unequal to the slope taken from the right-hand side. The function is continuous, and shading $x$ with $y$ generates a gradual pattern. (**D**) $y$ is a continuous and smooth function of $x$. At every point, the slope taken from the left-hand side is equal to the slope taken from the right-hand side (the slope at the maximal point being zero). A smooth function is always continuous. (**E**) With finite and noisy data, shading $x$ with $y$ often generates a bumpy pattern. It is unclear whether $y$ is a function of $x$, and if yes, whether the function is continuous (and perhaps even smooth).

#### 1.12 Considerations for selecting delay vector parameters for SSR

To construct delay vectors for SSR, one must choose the delay vector length $E$ and the time delay $\tau$. How does one choose $E$ and $\tau$? In general, detecting a continuous delay map requires that the delay vector length $E$ be high enough so that no two parts of the delay space cross. For example, using $E = 2$ (instead of $E = 3$) to make Figure 4C would have projected the delay space onto 2 dimensions. This would introduce line crossings, which would in turn produce artifactual discontinuities in the shading. On the other hand, the amount of data required to perform SSR inference is said to grow with the delay vector length [1]. SSR is less sensitive to $\tau$, although it is possible to mask a continuous delay map by choosing a "bad" $\tau$. For example, consider what would happen to Figure 4C if we set $\tau$ to the period of $Z$. Since the delay vector is $[Z(t), Z(t-\tau), Z(t-2\tau)]$, setting set $\tau$ to the period of $Z$ would force all 3 elements of the delay vector to always be equal. In geometric terms, this would compress the delay space onto a line, destroying the continuous delay map. However, bad choices of $\tau$ such as this are rare. In practice, a variety of methods are available for systematically choosing $E$ and $\tau$ [4, 5]. Appendix 1.13 discusses theoretical requirements for $E$ and $\tau$ in greater detail.

#### 1.13 Historical notes on the basis of SSR

SSR is about the existence and properties of delay maps. Here the term "map" is used interchangeably with "function": a map from $X$ to $Y$ sends every point in $X$ to one and only one point in $Y$. Different authors have focused on different aspects of delay maps when analyzing SSR, possibly introducing confusion over which aspect is the "correct" criterion for casuality. In particular, while CCM is thought to look for local smoothness of the map from the delay space of a causee to the values of a causer ([15]), [6] proposes that a method to detect continuity of the map is more consistent with the underlying theory. So is SSR about smoothness, or continuity? As we will see, Takens's theorem guarantees smoothness (a stronger criterion) but for a restricted set of cases, whereas the theorem of Sauer et al. essentially guarantees continuity (a weaker condition) but for a broader set of cases including fractals.

Takens's celebrated 1981 paper [9] is arguably the first major theoretical result underpinning a variety of data-driven methods for both causality detection (as discussed in this article) and forecasting (e.g. [7]). Theorem 1 of [9] is reproduced below, except that we have changed the names of some variables:

**Takens's theorem (theorem 1 of [9]):** Let $M$ be a compact manifold of dimension $m$. For pairs $(\phi, f)$, $\phi : M \to M$ a smooth diffeomorphism and $f : M \to \mathbb{R}$ a smooth function, it is a generic property that the map $\Phi_{(\phi,f)} : M \to \mathbb{R}^{2m+1}$, defined by

$$\Phi_{(\phi,f)}(p) = (f(p), f(\phi(p)), ..., f(\phi^{2m}(p))$$

is an embedding; by "smooth" we mean at least $C^2$.

We will attempt to illustrate Takens's theorem using the example in Figure S13. This system is described by a five-dimensional space of $[X, Y, Z, dX/dt, dY/dt]$ at various times (Figure S13A). However, once we have chosen the five initial conditions, $X$ and $Y$ uniquely determine $dX/dt$ and $dY/dt$. Thus we visualize the state space in just three dimensions $(X, Y, Z)$ (Figure S13C), and color the trajectory with time (a colored ring in Figure S13D, similar to a clock to highlight the periodic nature of system dynamics). This trajectory is the manifold $M$ in Takens's theorem and is 1-dimensional ($m = 1$) since it is a loop. We can think of $\phi$ as a function that maps a point $p$ on the manifold $M$ at the current time $t$ to the point $q$ at a a previous time $t - \tau$, and therefore $\phi^2(p)$ would operate $\phi$ twice and map $p$ at the current time $t$ to the point $r$ at time $t - 2\tau$ (olive in Figure S13B) and $\phi^{-1}(q)$ would map point $q$ at the past time $t - \tau$ to the point $p$ at the current time $t$ (Figure S13C). The term "diffeomorphism" in the theorem means that both this function $\phi$ and its inverse function (the map from past to present) are smooth (Figure S12). $f$ can be viewed as an "observation" function that maps each point on the manifold to a single real number (e.g. in Figure S13E, $f(p) = p_Z$ so that $f$ simply returns the $Z$ coordinate of point $p$). Takens's theorem then asks us to consider a function $\Phi$ that maps a point $p$ at time $t$ on our state space manifold (Figure S13E) to a point in the "delay space" with coordinates consisting of the values of the observation function applied to point $p$ at time $t$, point $q$ at time $t-\tau$, and point $r$ at time $t-2\tau$ (e.g. $[p_Z, q_Z, r_Z]$ in Figure S13F). This choice of delay space comes from three earlier choices: First, we consider delayed values of $Z$ since $Z$ is what our observation function $f$ returns; second, since $m = 1$, the delay space should be of dimension 3 ($= 2m+1$); third, the delay $\tau$ comes from our diffeomorphism $\phi$. Then, Takens's theorem states that for "most" (technically, "generic") choices of $f$ and $\phi$, $\Phi$ is an embedding. This means that $\Phi$ is diffeomorphic to its image, i.e. the curve in delay space will map smoothly to the state space manifold and vice versa [12].

Indeed from Figure S13 (C-F), we can see that when the observation function $f = Z(t)$, $\Phi$ from the state space to the delay space is a map. This is because each dot in the state space corresponds to a single time color (i.e. a point within a period), and each time color corresponds to a single dot in delay space, and thus, each point in the state space corresponds to a single point in the delay space. Moreover, $\Phi$ is continuous because the maps from state space to the time ring and from the time ring to delay space are both continuous. Similarly, the inverse of $\Phi$ from delay space to state space is also a continuous map. Moreover, we know from solutions to our equations that both maps are additionally smooth. Thus, we have verified Takens's theorem for this observation function.

Strikingly, if the observation function is $Y$, we will no longer have a continuous map from the delay space of $Y$ to the state space. This is visualized as "bumpy coloring" in Figure S13H. In fact, we cannot even map the delay space to the time ring or the state space: $(p, q, r)$ and $(p', q', r')$ occupy the same point in the $Y$ delay space, yet correspond to different times within a period (Figure S13B) and thus they correspond to different locations in the state space. In summary, we cannot map the $Y$ delay space to state space. Takens's theorem took care of this pathology using the word "generic". That is, $Y$ is not considered a generic observation function here. On the other hand, if we use an observation function based on 95% $Y$ mixed with 5% $Z$, we get an embedding from the state space to the delay space (Figure S13I-J). This is essentially what the term "generic" means in the context of topology: Although some observation functions do not give you an embedding, these "bad" observation functions can be tweaked just a little bit to become "good" ones. Similarly, some choices of $\phi$ do not work (i.e. $\tau = T$ for this system), but these are exceptions (see [12] Theorem 2 for what makes a $\phi$ "generic").

Roughly speaking, Sugihara and colleagues essentially used the values of one variable to shade the delay space of another variable, and used the local smoothness of the map to infer causality. In the example of Figure 4 in the main text, shading delay space of $Z$ with $Y$ generates a continuous (and smooth) pattern, consistent with $Y$ causing $Z$. On the other hand, shading delay space of $Z$ with $W$ shows a bumpy pattern, consistent with $W$ not causing $Z$.

Sauer and colleagues [11] later extended this work by proving a similar result that is in some ways more general. Theorem 2.5 in [11] is distinct but related to Takens's theorem, and applies to cases that Takens's theorem does not cover, such as fractal spaces. Additionally, [11] replaces the concept of "generic" with a different notion ("prevalence"), which is closer to a probabilistic statement. Cummins et al. then proved an important corollary to Sauer's result (corollary 4.3 in [6]), which showed that even under the more general conditions of [11], the map from the delay space to the state space is continuous.

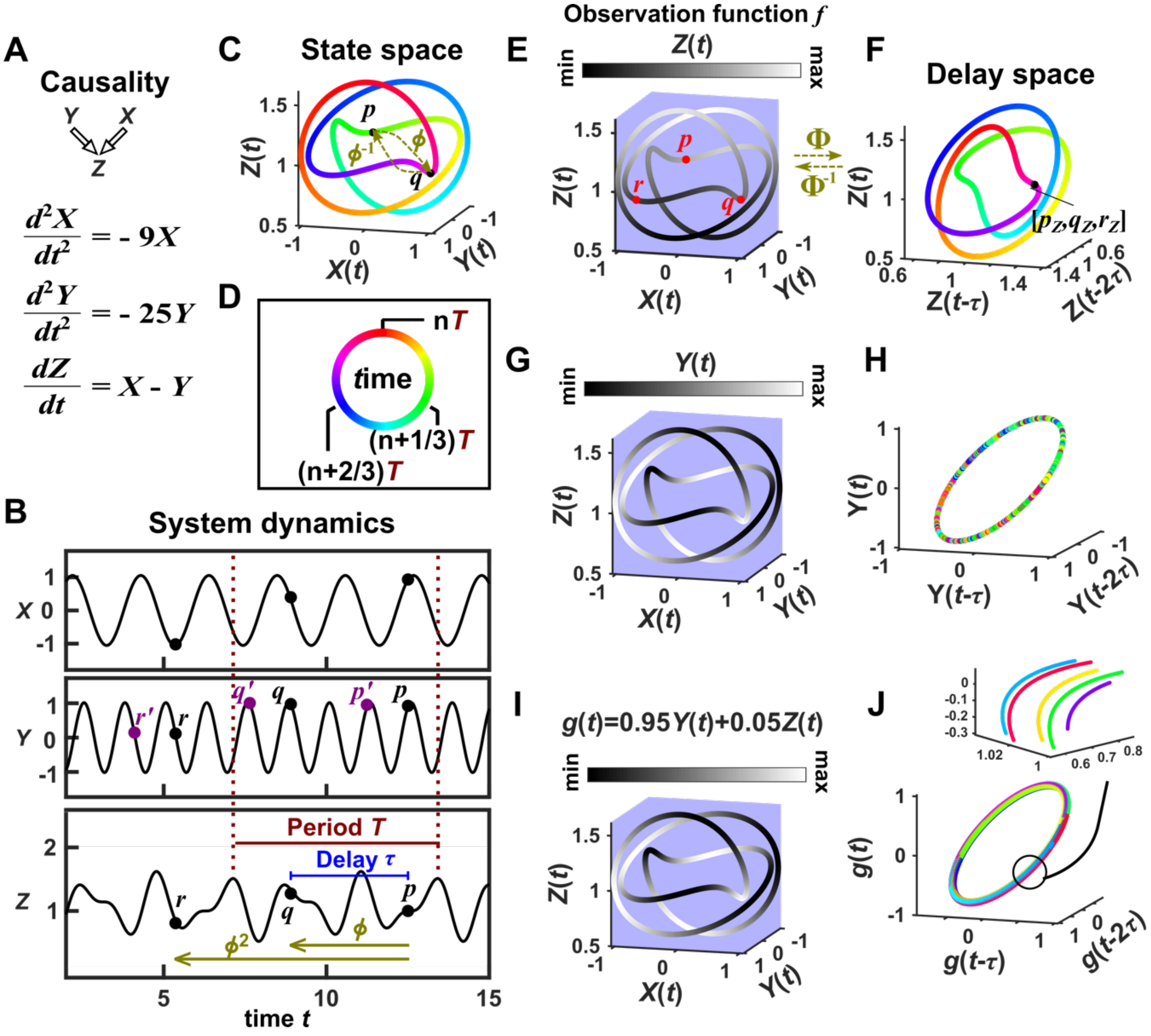


Figure S13: Illustration of Takens's theorem. (**A**) We consider a 3-variable toy system in which $X$ and $Y$ causally influence $Z$, but $Z$ does not influence $X$ or $Y$. (**B**) Time series of the three variables. (**C**) We can plot time series data in the state space $M$. Takens's theorem requires that $\phi$, a function that maps a point $p$ at current time $t$ to the point $q$ at a previous time $t-\tau$, and its inverse $\phi^{-1}$ (from past to current) are both smooth ($C^2$: the first and second derivatives of the function exist and are continuous for all time). To mark time progression, we color each point along the trajectory with its corresponding time value where time is represented as a color ring similar to a clock to reflect the periodic nature of system dynamics (**D**). (**E, G, I**) Shading the state space with the observation function ($f$ in Takens's theorem) marked above. (**F, H, J**) Delay space based on the observation function, colored with time. The map $\Phi$ in Takens's theorem maps, for example, point $p$ in E to point $[p_Z, q_Z, r_Z]$ in F. The theorem states that for "generic" observation functions, this map $\Phi$ and its inverse $\Phi^{-1}$ are both smooth (differentiable). In this example $\tau = 3.6$. In J, multiple colors in a region are due to one period wrapping around the delay space multiple times (inset), but the color shading transition is gradual (similar to F).

#### 1.14 Nonreverting continuous dynamics: Criteria and effects on convergent cross mapping

We first illustrate "nonreverting continuous dynamics", which is SSR's version of nonstationarity. We then discuss how nonreverting continuous dynamics affects CCM.

We use the phrase "nonreverting continuous dynamics" to describe the following idea: If the $X$ delay space maps continuously to $t$, and $t$ maps continuously to $Y(t)$, then $X$ delay space will map continuously to $Y(t)$, even if $X$ and $Y$ are causally unrelated (Figure S14A). Figure S14B illustrates this with three causally *independent* time series $X$ and $Y$. In the top row, the $X$ delay space maps continuously to $t$ ("nonreverting") and $t$ maps continuously to $Y(t)$ ("continuous"), so we get nonreverting continuous dynamics and a continuous delay map from $X$ to $Y$ even though $X$ and $Y$ are independent. In the middle row, the $X$ delay space maps continuously to $t$ , but $t$ does not map continuously to $Y(t)$, so we do not have nonreverting continuous dynamics, nor a continuous map from the $X$ delay space to $Y$. In the bottom row, the $X$ delay space does not map continuously to $t$. This is because a single delay vector (represented by, for example, cyan segments) maps to many time points, generating a bumpy pattern similar to Figure S12A. We refer to this lack of a continuous map from delay space to time as "reverting". In this case, even though $t$ maps continuously to $Y(t)$, we do not have nonreverting continuous dynamics and we do not get a spurious continuous map from the $X$ delay space to $Y$.

While our visual inspection method is useful for diagnosing this pathology for noiseless data in low-dimension delay space, visualization becomes difficult in $> 3$ dimensions and is complicated by noise. A simple idea to check for this problem in real data might be to quantify cross map skills from the $X$ delay space to $t$ and from $t$ to $Y$. If both cross map skills are high, then we might have nonreverting continuous dynamics. In this case, high cross map skill from $X$ delay space to $Y$ may not suggest that $Y$ causes $X$.

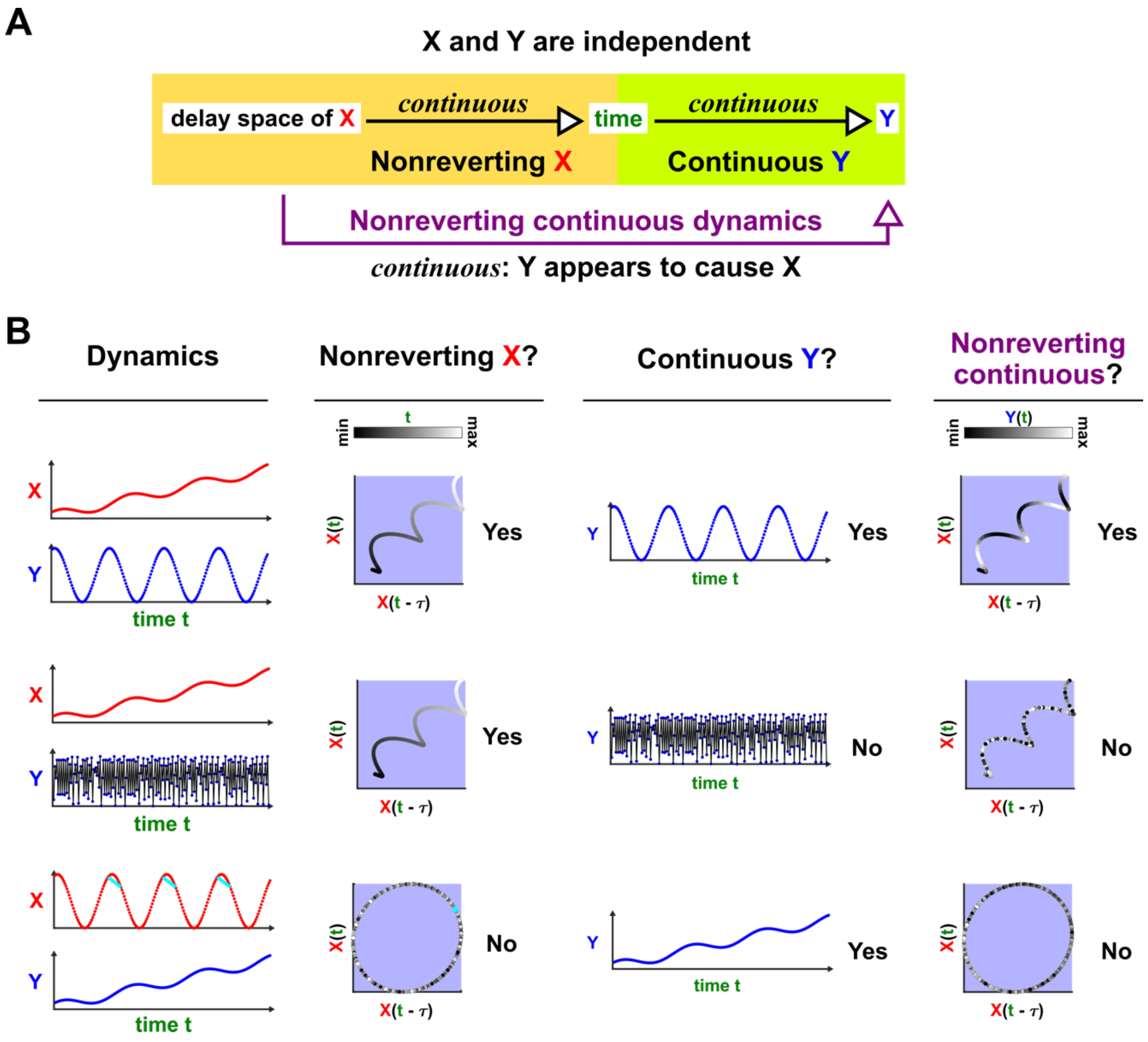


Figure S14: Nonreverting continuous dynamics. (**A**) Nonreverting continuous dynamics. We call $X$ reverting if the delay space of $X$ maps continuously to time. We call $Y$ "continuous" if $Y(t)$ is continuous with respect to time (i.e. time maps continuously to $Y(t)$). If $X$ is reverting and $Y$ is continuous then we say that the pair of time series $X, Y$ has nonreverting continuous dynamics. (**B**) Examples. In each row, $X$ and $Y$ are causally independent. Leftmost column: Dynamics. Each red or blue dot represents a single time point visible upon zooming in. Second column: Testing for a continuous map from the delay vectors of $X$ ($X$ delay space) to $t$ (time), i.e. nonreverting $X$ dynamics. Third column: Testing for a continuous map from $t$ to $Y$ (continuous $Y$ dynamics: whether $Y$ at nearby time points share similar values). Since the data points are discrete, $Y$ cannot be truly continuous in the mathematical sense, so when dealing with real data, "continuous $Y$" really means "highly autocorrelated". Fourth Column: the presence or absence of "nonreverting continuous dynamics". With nonreverting continuous dynamics, since there is a continuous map from $X$ delay space to $Y$, $Y$ appears to cause $X$ even though $X$ and $Y$ are causally independent.

Nonreverting continuous dynamics interferes with CCM causal inference. In particular, when quantifying cross map skill, one could intersperse training and testing data when the system is nonstationary [10] (Figure S15, Column 4). However, we find that this approach leads to false positive errors (Figure S15 bottom row). In contrast, the alternative (not interspersing training and testing data) leads to false negative errors (Figure

S15, third row). Thus, the ability to correctly infer causality with CCM is vastly reduced when data exhibit nonreverting continuous dynamics.

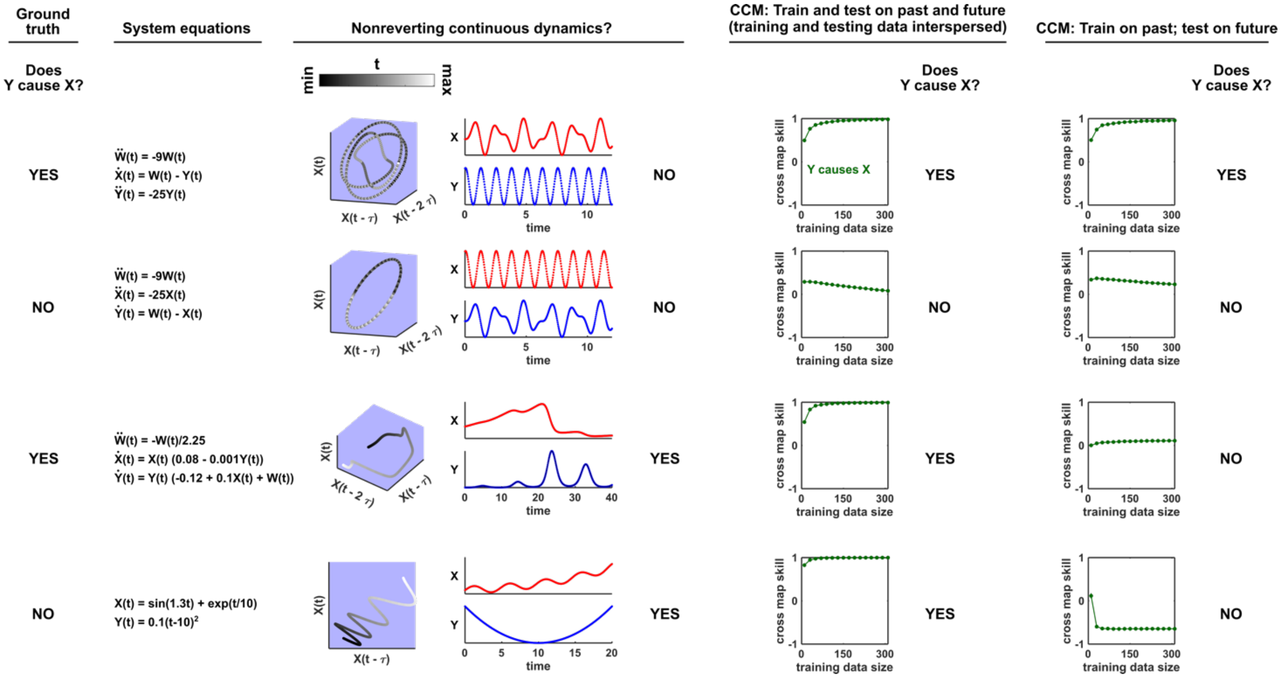


Figure S15: With nonreverting continuous dynamics, CCM results may depend on analysis details (i.e. the choice of training and testing data) instead of ground truth. Each row represents a system where $Y$ may or may not causally influence $X$ (Column 1). Column 2: Governing equations. Column 3: Checking for nonreverting continuous dynamics as in Figure S14. The top two rows do not have nonreverting continuous dynamics since there is no continuous map from the delay space of $X$ to time. The bottom two rows have nonreverting continuous dynamics. Columns 4 and 5: Results of CCM where training and testing data are interspersed or when we train on the past and test on future.

#### 1.15 Possible failure modes of the prediction lag test

State space reconstruction methods suffer false positive errors in the presence of synchony [1]. This occurs when "the dependence of the dynamics of the forced variable on its own state is no longer significant" [1]. Ye et al. proposed a test in an effort to solve this problem [3]. Their procedure relies on finding mappings from the delay vector $[X(t), X(t-\tau), X(t-2\tau)...X(t-(\text{E-1})\tau)]$ to $Y(t+l)$, where $E$ is the delay vector length, $\tau$ is the time lag, and $l$ is a key variable known as the "prediction lag". They then examine how the cross map skill (Figure 6B) varies with the prediction lag. According to this technique, if the cross map skill is maximized at a positive prediction lag ($l > 0$), then the putative causality is spurious and arose from, for example, strong unidirectional forcing. On the other hand, if the highest quality mapping occurs at a non-positive prediction lag ($l \leq 0$), then we have further evidence that the detected causality is real and not spurious.

We find that while this test correctly distinguishes between real and spurious causal signals at some times, at other times it does not. In Figure S16, we explore the behavior of this test. Within each row of Figure S16, we examine a different system and ask whether $Y$ causes $X$ according to: (1) the ground truth model, (2) our visual continuity test, (3) a CCM cross map skill test (without the prediction lag test), and (4) the prediction lag test.

In rows 1 and 2 of Figure S16, the prediction lag test performs well, overturning the results of the visual continuity and CCM tests when apparent causality is spurious (row 1), and agreeing with the continuity and CCM tests (row 2) when apparent causality is real (modified from [3] Eq. 2). However in row 3, the prediction lag test dismisses a true causal link as spurious. Moreover, when we apply the prediction lag test to a system with a periodic putative driver (Figure S16 row 4), we find that cross map skill is a periodic function of the prediction lag. While this result is precisely what we would expect mathematically, its causal interpretation is unclear. The fifth row of Figure S16 takes the idea of strong forcing to the extreme, so that $Y(t+1)$ is a function of $X(t)$, but not $Y(t)$. Here the prediction lag test gives a false positive error. In the bottom row, $X$ and $Y$ do not interact, but are both driven by a common cause $W$ with different lags. Specifically, $W(t)$ exerts a direct effect on $Y(t+1)$ and on $X(t+3)$. Thus, $Y$ receives the same information as $X$, but at an earlier time, analogous to Figure 3iii. Consistent with this, delay vectors of $X$ predict past values of $Y$ better than future values of $Y$. Thus, the prediction lag test produces a false positive error.

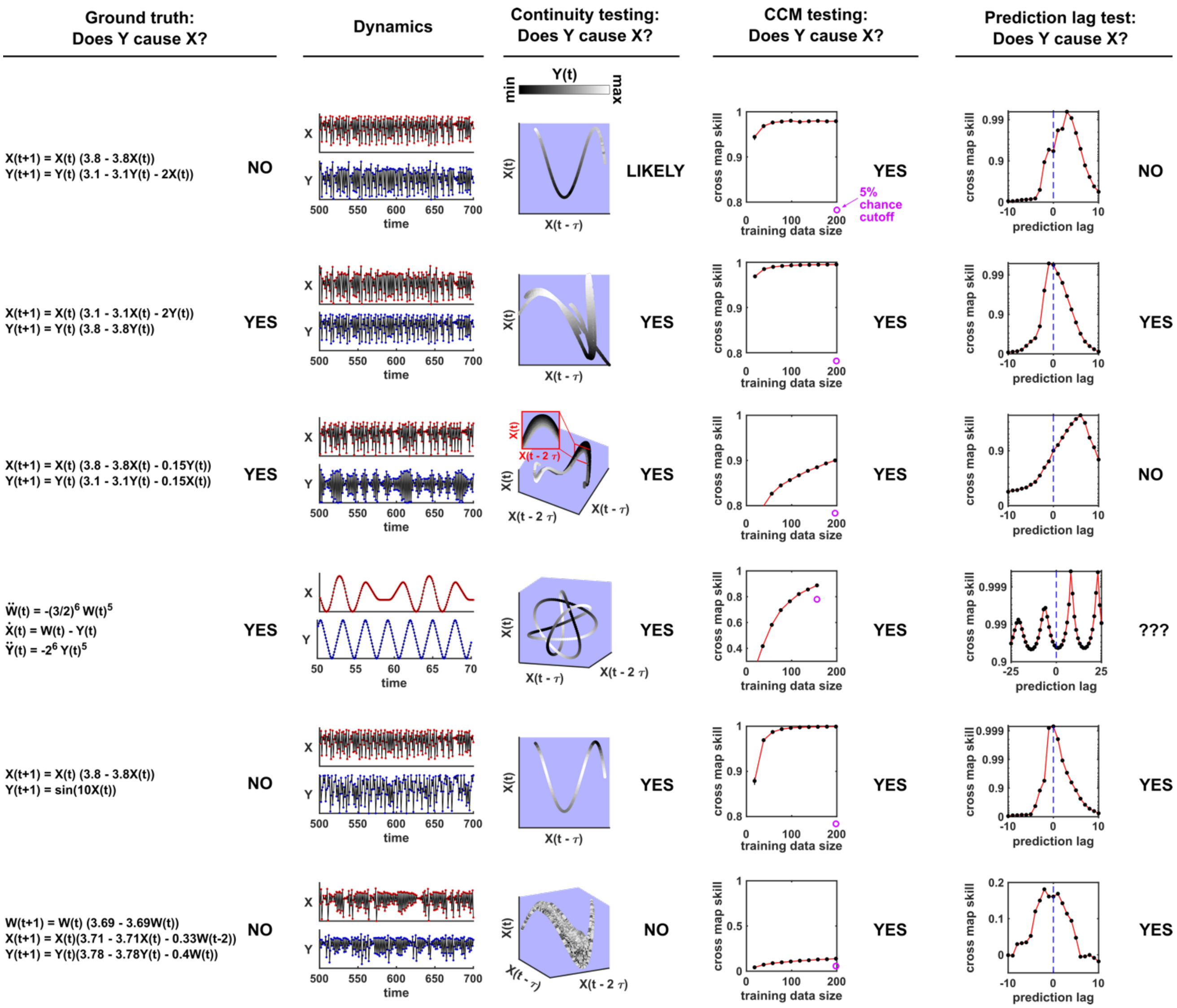


Figure S16: Comparison of visual continuity testing, cross map skill testing, and prediction lag testing in causal inference. Each row represents a two-variable or three-variable system where $Y$ may or may not causally influence $X$. The leftmost column shows the equations and ground truth causality. The second column shows a sample of $X$ and $Y$ dynamics. Red and blue dots represent $X$ and $Y$ values, respectively; black lines connecting the dots serve as a visual aid. The third column shows visual continuity testing and causal interpretation. We write "likely" in the top row because the map from $X$ delay space to $Y$ appears to have some small bumps on the right side of the plot. The fourth column shows cross map skill testing (without the prediction lag test) and causal interpretation. Black dots show cross map skill. Open purple dots show the 5% chance cutoff at the maximum library size according to random phase surrogate data testing (see Supplementary Methods), or are placed below the horizontal axis if the 5% chance cutoff is below the plot. In all systems $Y$ appears to cause $X$ according to cross map skill testing since cross map skill is positive, increases with training data size, and is significant according to the surrogate data test. The rightmost column shows the prediction lag test and causal interpretation.

Ye et al. [3] applied the prediction lag test to 500 systems with the same form as in the third row of Figure S16 but with randomly chosen parameters. They found that within the parameter range they sampled, false negative errors as in Figure S16 do occur, but such errors are rare. We repeated the randomized numerical experiment from [3] for both the original parameter range of [3] (Figure S17B, "friendly" parameter regime) and a second parameter range of the same volume in parameter space (Figure S17B, "pathological" parameter regime). In this pathological parameter regime, false negative errors occur in the overwhelming majority of cases.

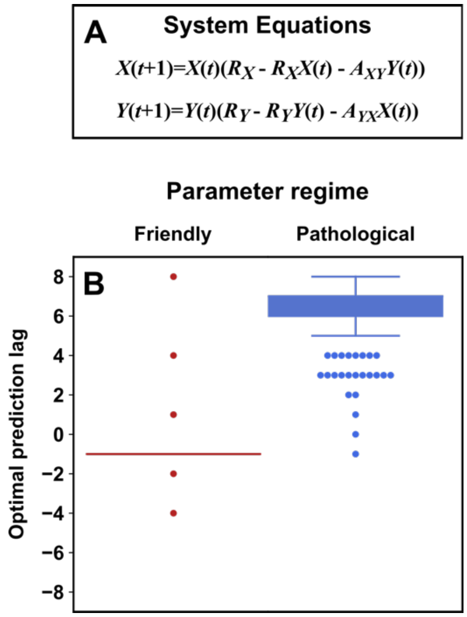


Figure S17: Parameters within a "pathological" regime almost always cause the prediction lag test of [3] to erroneously reject a true causal link. (**A**) System equations. For both "friendly" and "pathological" regimes, initial conditions $X(1)$ and $Y(1)$ were independently and randomly drawn from the range $0.01-0.99$, and $R_X$ was randomly drawn from the range $3.7-3.9$. $R_Y$ was randomly drawn from the range $3.7-3.9$ ("friendly") or $3.1-3.3$ ("pathological"). $A_{XY}$ and $A_{YX}$ were independently (and randomly) drawn from the range $0.05-0.1$ ("friendly") or $0.15-0.2$ ("pathological"). (**B**) Boxplots show the optimal prediction lag when using delay vectors made from $X$ to predict values of $Y$ in 500 systems with randomly selected parameters. In the ground truth model for this system, $Y$ exerts a causal influence on $X$. In the "friendly" parameter regime explored in [3], the optimal prediction horizon is negative, correctly indicating that $Y$ does indeed cause $X$. In the "pathological" regime, the optimal prediction horizon is positive, and so the rule of [3] would wrongly lead us to conclude that $Y$ does not cause $X$. In the friendly regime the "box" is shown as a line because the vast majority of trials had the same optimal prediction lag of $-1$.

### 2 Supplementary Methods

#### Methodological details for Figure 2

For panel B, we simulated the random walk system

$$X(t+1) = X(t) + \epsilon(t)$$

where $\epsilon(t)$ terms were drawn independently from a normal distribution with mean of 0 and standard deviation of 1. We simulated this system from the initial condition of $X(1) = 0$ through 999 subsequent steps. For panel C, we simulated the autoregressive system

$$X(t+1) = 0.75X(t) + 10 + \epsilon(t)$$

where $\epsilon(t)$ terms were again drawn independently from a normal distribution with mean of 0 and standard deviation of 1. We simulated this system from the initial condition of $X(1) = 40$ for 1999 subsequent steps. We only used the final 1000 steps for computing the correlation between two time series.

To compute the significance of the Pearson correlation between two time series, we used surrogate data generated by either permutation or the random phase procedure. Permutation surrogate time series were generated by randomly shuffling data. Random phase surrogate time series were generated by Ebisuzaki's random phase method [13] as implemented in the rEDM (version 1.5) function make_surrogate_data. For a pair of time series $[X_1(1), X_1(2), ..., X_1(1000)], [X_2(1), X_2(2), ..., X_2(1000)]$, we first computed the Pearson correlation $\hat{\rho}$ between the two time series. We then replaced the $X_2$ values with surrogate time series and recomputed the Pearson correlation as $\tilde{\rho}$. We computed this shuffled correlation 9999 times (permutation) or 499 times (random phase) to get a null distribution $[\tilde{\rho}_1, \tilde{\rho}_2, ..., \tilde{\rho}_n]$. Following [18], we computed the $p$ value as

$$p = (N_{stronger} + 1)/(N_{surr} + 1) \tag{12}$$

where $N_{surr}$ is the number of surrogates, $N_{stronger}$ is the number of surrogate correlations $\tilde{\rho}$ whose magnitude was greater than or equal to the magnitude of the original correlation $\hat{\rho}$, and the "+1" terms account for $\hat{\rho}$.

#### Methodological details for Figure 4

The system of equations was numerically integrated using the ode45 method in Matlab from $t = 0$ to $t = 200$ in time steps of 0.03, and plotted in the delay space $Z$ with $\tau = 3.6$. The initial condition for all state variables ($V$, $W$, $X$, $Y$, $Z$, $\frac{dX}{dt}$, $\frac{dY}{dt}$, and $\frac{dV}{dt}$) was 1. For panel F, measurement noise was added to $Y(t)$ . Specifically, noisy data were generated as:

$$Y^{obs}(t) \sim \text{Unif}\left(Y(t) - 3^{1/2}\left(0.15\Delta_Y\right), Y(t) + 3^{1/2}\left(0.15\Delta_Y\right)\right)$$

where Unif$(a, b)$ is a uniform random variable bounded by $a$ and $b$, and $\Delta_Y$ is the difference between the maximum and minimum values of $Y(t)$ between $t = 0$ and $t = 200$. These noise parameters are chosen so that $Y^{obs}(t)$ is centered at $Y(t)$ and has a standard deviation of $0.15\Delta_Y$ .

#### Methodological details for Figure 5

The dynamics in the top row of Figure 5 were generated from the equations:

$$X(t) = \sin(t) + 0.5t$$
$$Z(t) = 0.1(t - 10)^2$$

This continuous-time system was discretized from $t = 1$ to $t = 20$ on an evenly spaced grid of 400 data points for visualizing delay spaces where the time delay is 50 time points (i.e. $\tau = 50(20 - 1)/(400 - 1)$).

The dynamics in the second row of Figure 5 were generated from the equations:

$$X(t + 1) = X(t)(3.61 - 3.61X(t))$$
$$Z(t + 1) = Z(t)(3.61 - 3.61X(t))$$

with initial conditions of $X(1) = 0.4$ and $Z(1) = 0.7$. For this system, $\tau = 1$ and $t = 1, 2, ..., 2000$ were used to make the plots of delay spaces.

The dynamics in the third row of Figure 5 were generated from the equations:

$$\frac{dX^2}{dt} = -X(t)$$
$$\frac{dZ^2}{dt} = -25Z(t)$$

with initial conditions of $X(1) = X'(1) = Z(1) = Z'(1) = 1$. For this system, $\tau = 0.9$ was used for delay spaces. This continuous-time system was numerically integrated using the ode45 method in Matlab from $t = 0$ to $t = 13.998$ on a grid of 4667 evenly-spaced time points for plotting dynamics, and time points $t = 0.003$ through $t = 7.698$ were used for visualizing delay spaces.

The dynamics in the bottom row of Figure 5 were generated from the classic Lorenz attractor equations:

$$\frac{dX}{dt} = -10X(t) + 10Y(t)$$
$$\frac{dY}{dt} = 28X(t) - Y(t) - X(t)Z(t)$$
$$\frac{dZ}{dt} = -\frac{8}{3}Z(t) + X(t)Y(t)$$

with initial conditions of $X(0) = Y(0) = Z(0) = 1$. A delay of $\tau = 0.14$ was used to make delay spaces. This continuous-time system was numerically integrated using the ode45 method in Matlab from $t = 0$ to $t = 399.98$ on an evenly spaced grid of 5715 data points for visualizing delay spaces.

### Methodological details for Figure 7

#### Ground truth model and data generation

We used the ground truth model:

$$S_1(t+1) = \max\left(0, S_1(t)\left(1.2 - 0.1S_1(t) + D_1(t)\right) + \epsilon_{p1}(t)\right)$$
$$S_2(t+1) = \max\left(0, S_2(t)\left(1.1 - 0.2S_2(t) + D_2(t) + 0.3S_1(t)\right) + 2.5\epsilon_{p2}(t)\right)$$

$S_1(t)$ and $S_2(t)$ represent the population sizes of species 1 and 2 at time $t$. $D_1(t)$ and $D_2(t)$ are the values of periodic drivers at time $t$. Specifically, in both the two-driver and one-driver cases:

$$D_1(t) = 0.05\sin\left(t + \phi_1\right) + 0.05\sin\left(\frac{5t}{6} + \phi_1\right)$$

In the two-driver case:

$$D_2(t) = 0.1\sin\left(\frac{t}{\sqrt{10}} + \phi_2\right)$$

Conversely, in the one-driver case $D_2(t) = 0$. The process noise terms $\epsilon_{p1}(t)$ and $\epsilon_{p2}(t)$ are both IID normal random variables with mean of 0 and with shared standard deviation $\sigma_p$. Specifically, for any pair of times $t_1 \neq t_2$, $\epsilon_{p1}(t_1)$ and $\epsilon_{p1}(t_2)$ are independent, and similarly for $\epsilon_{p2}$. Also, all values $\epsilon_{p1}(1), \epsilon_{p1}(2), \cdots$ are independent of all values $\epsilon_{p2}(1), \epsilon_{p2}(2), \cdots$. At the beginning of each replicate simulation, the phases $\phi_1$ and $\phi_2$ are independently assigned a random number from a uniform distribution between 0 and $2\pi$, and do not change with time.

To generate data without measurement noise, we simulated this system for $t = 1, 2, ..., 400$ with the initial conditions $S_1(1) = 2$; $S_2(1) = 4.5$. We used the final 200 time points for inference to help ensure that the system had reached equilibrium behavior.

We also introduced additive measurement noise to simulate instrument uncertainty:

$$S_1^{obs}(t) = S_1(t) + \epsilon_{m1}(t)/1.5$$
$$S_2^{obs}(t) = S_2(t) + \epsilon_{m2}(t)$$

where $S_1^{obs}$ and $S_2^{obs}$ represent the observed values (i.e. noisy measurements) of $S_1$ and $S_2$. $\epsilon_{m1}(t)$ and $\epsilon_{m2}(t)$ are also IID normal random variables with mean of 0 and standard deviation $\sigma_m$. The tables in Figure 7D are generated by varying $\sigma_p$ from 0 to 8 and varying $\sigma_m$ from 0 to 1.

#### Causal inference using Granger causality and CCM

For each combination of $\sigma_m$ and $\sigma_p$ (the standard deviation of measurement noise and process noise, respectively), we generated 1000 time series for $S_1$ and $S_2$ as described above. For each replicate pair of time series, we used Granger causality and CCM to infer whether $S_1$ causes $S_2$ (it does) and whether $S_2$ causes $S_1$ (it does not).

#### Granger causality inference

We used the multivariate Granger causality Matlab package (MVGC, [14]). We used the following settings:

- regmode = 'OLS' (We fit the autoregressive model by the ordinary least squares method).
- icregmode = 'LWR' (We determined the information criterion using the LWR algorithm. This is the default setting).
- morder = 'AIC' (We used Akaike information criterion to determine the number of lags in the autoregressive model).
- momax = 50 (We used a maximum of 50 lags in the autoregressive model).
- tstat = ” (We used Granger's F-test for statistical significance. This is the default setting).

We inferred the presence of a causal link if the p-value was less than or equal to 0.05. We inferred no causal link otherwise. When $\sigma_m$ and $\sigma_p$ were both 0, the MVGC package (correctly) exited with an error on most trials. We reported this as "unsuitable data" in Figure 7D & E.

When $\sigma_m$ and $\sigma_p$ are both 0, the inferred spectral radius of the stochastic process is close to 1, and the MVGC routines can be prohibitively slow (i.e. when running 1000 trials, the program would hang at an early stage for hours). In this case, the authors note that switching from the package's default single-regression mode to an alternative dual-regression mode may improve runtime [14]. We thus switched to the dual-regression mode when the spectral radius was between 0.9999 and 1 (a spectral radius of 1 or more causes an error). This fix had no effect on benchmark results as long as at least one of $\sigma_m$ and $\sigma_p$ was not 0.

##### Convergent cross mapping

Convergent cross mapping looks for a delay map from $X$ to $Y$. That is, CCM looks for a map from $[X(t), X(t-\tau), X(t-2\tau), ..., X(t-(E-1)\tau)]$ to $Y(t)$. Thus in order to apply CCM one needs to choose the delay $\tau$ and the vector length (dimension of the delay space) $E$. $E$ and $\tau$ should ideally be "generic" in the sense of Takens's theorem: we want to avoid line-crossing (such as the symbol "$\infty$") in the delay space, because otherwise, $\Phi^{-1}$ in Figure S13 does not exist. There are different ways to do this, but no method is obviously the best ([4, 5]).

Following [4] and [1] we chose $\tau$ and $E$ to maximize univariate one-step-ahead forecast of the putative causee $X$. That is, for $X(n)$, we try to predict $X(n+1)$ using the simplex projection method by finding delay vectors in the training data of $X$ that are most similar to $[X(n), X(n-\tau), X(n-2\tau), ..., X(n-(E-1)\tau)]$, and take weighed average of their $X$ values 1 step in the future (i.e. Figure 6A where $X = Y$ and the prediction lag is 1). If the delay space has a line crossing, then at the cross-point, a one-step-ahead forecast may have more than one possible outcome and thus perform poorly. In more detail, we made one-step-ahead forecasts within the time range 201-400 (we did not use time range 1-200 to avoid transient dynamics). As per the field standard, we used leave-one-out cross-validation to do simplex projection. That is, when making a forecast for a time $t$, we used all times within 201-400 other than $t$ as training data (200 time points). We performed a grid search, varying $\tau$ from 1 to 6 and varying $E$ from 1 to 6. We then used the combination of $\tau$ and $E$ that maximized the forecast skill (the Pearson correlation between forecasts and true values) for subsequent CCM analysis. Additionally, following [1], if the optimal combination of $\tau$ and $E$ failed to give a significantly positive forecast skill, we did not report CCM results for that trial and reported the trial as "unsuitable data". To test whether forecast skill is "significantly positive", we ask whether it is robust to small changes in the training dataset. To do so, we used a naive bootstrap approach to create different versions of training libraries composed of randomly chosen delay vectors (sampling with replacement: some vectors may not be sampled and others may be sampled more than once) from the original training data using the 'random_libs' setting in the rEDM (version 1.5) ccm method. The training library size (the number of delay vectors in the library) was chosen to be 200. We then calculated forecast skills with 300 such libraries and considered the forecast skill "significant" if at least least 95% gave a forecast skill greater than 0.

Having chosen $\tau$ and $E$, we used three CCM criteria to infer causality (criteria 1-3 in Figure 6). We did not use the fourth criterion (the prediction lag test) since its interpretation is unclear for periodic systems (Figure S16). For all three criteria, we used the same cross-validation setting that we used to choose $\tau$ and $E$. The first CCM criterion is that cross map skill is greater than 0. Thus, we computed cross map skill using the maximum possible number of distinct delay vectors ($200-(E-1)\tau$) and compared this value to 0.

The second CCM criterion is that the cross map skill from causee to causer with real data must be greater than the cross map skill when the putative causer is replaced with surrogate data. To test this criterion, we first computed cross map skill using the same training and testing time points as before to obtain a single cross map skill value. We then repeatedly (1000 times) computed cross map skill in the same way, but now with the putative causer time series replaced with random phase surrogate data. Random phase surrogate data were generated by Ebisuzaki's method as implemented in the rEDM function make_surrogate_data. We then computed the $p$-value according to Eq. 12. A putative causal link would pass this criterion if the $p$-value was less than or equal to 0.05.

The third CCM criterion is that cross map skill increases with more training data. Following [4], we again used a naive bootstrap approach to test for this criterion. Specifically, we computed the cross map skill with a training library composed of randomly chosen delay vectors sampled with replacement from the original training data time points. We used either a large library with $200-(E-1)\tau$ available training vectors as used previously, or a small library with 15 training vectors. For each of 1000 bootstrap trials, we compared the cross map skill from a randomly chosen small library to the cross map skill from a randomly chosen large library. We said that the cross map skill increased with training data if the cross map skill of the large library was greater than that of the small library in at least 95% of the 1000 bootstrap trials.

For "alternative" CCM testing, we only changed how the third CCM criterion (cross map skill increases with more training data) were tested. Here, instead of using the bootstrap test of [4], we tested the third CCM criterion using Kendall's $\tau$ test as suggested in [16]. To do this, we varied the library size from a minimum of 15 vectors to the the maximum library size ($200-(E-1)\tau$), in increments of 3 vectors. For each library size, we computed cross map skill using 50 libraries randomly sampled without replacement (e.g. the 50 libraries would be identical at the maximal library size). We then computed the median cross map skill for each library size. Finally we ran a 1-tailed Kendall's $\tau$ test for a positive association between library size and median cross map skill. We used the function stats.kendalltau from the Python package SciPy to compute a 2-tailed $p$-value, and then divided this $p$-value by 2 to get a 1-tailed $p$-value. We said that cross map skill increased with training data if the $\tau$ statistic was positive and the 1-tailed $p$-value was $\leq 0.05$.

#### Methodological details for Figure S3

The original subpopulation distributions are normal distributions with standard deviation of 10 and mean of 100 (male) or 130 (female). Each sampling plot shows 300 samples randomly drawn from the appropriate mixture distribution.

#### Methodological details for Figure S13

To generate data for panels C-J, the system of panel A was numerically integrated using the ode45 method in Matlab with a time step of 0.005 and with the initial condition that $X$, $Y$, $Z$, $\frac{dX}{dt}$, $\frac{dY}{dt}$ were all set to 1 at $t = 0$. Panels C , E, F, G, I, and J (non-inset) show data from a single period. For panel H the system was integrated for about 5 periods to more clearly visualize the lack of a continuous delay map. For the panel J inset the system was integrated for about 12 periods to better see the separated legs of the curve upon zooming in. Panels C, D, F, H, and J were colored $\text{mod}(t, T)$. That is, they were colored by the remainder of $t$ (time) after dividing by $T$ (here $T = 2\pi$). $\tau = 3.6$ was used for all delay spaces.

#### Methodological details for Figure S14

All systems were discretized from $t = 1$ to $t = 20$ on an evenly spaced grid of 200 points for visualizing delay spaces.

The dynamics in the top row were generated from the equations:

$$X(t) = \sin(t) + 0.5t$$
$$Y(t) = \sin(1.3t)$$

A delay time of 12 time indices (i.e. $\tau = 12(20-1)/(200-1)$) was used for constructing delay spaces.

The dynamics in the second row were generated from the equations:

$$X(t) = \sin(1.3t)$$
$$Y(t) = Y(t-\delta)(3.77 - 3.77 * Y(t-\delta))$$

with $\delta = (20-1)/(200-1)$ and the initial condition $Y(1) = 0.3$. A delay time of 25 time indices (i.e. $\tau = 25(20-1)/(200-1)$) was used for constructing delay spaces.

In the third row the dynamics are identical to the first row, except that $X$ and $Y$ are switched, and $\tau = 25$ time indices was used for constructing delay spaces.

#### Methodological details for Figure S15

Top row: For this system, we used the initial conditions $W(0) = \dot{W}(0) = X(0) = Y(0) = \dot{Y}(0) = 1$. We numerically integrated this system using ode45 in Matlab with a time step of 0.03. We composed delay vectors of length $E = 3$ with a delay of $\tau = 3.6$. We visualized the delay space using data from $t = 0$ through $t = 29.97$ (time indices 1-1000). For CCM with temporally separate training and testing sets, we used data from $t = 0$ through $t = 14.97$ (time indices 1-500) for training data and data from $t = 15$ through $t = 29.97$ (time indices 501-1000) for testing. Specifically, in the rEDM (version 0.7.2) ccm method we set the lib argument to "(1, 500)" and set the pred argument to "(501, 1000)". For CCM with temporally interspersed training and testing sets, we set both the lib and pred arguments to "(1,1000)". This setting instructs rEDM to use leave-one-out cross-validation.

Second row: For this system, we used the initial conditions $W(0) = \dot{W}(0) = X(0) = \dot{X}(0) = Y(0) = 1$. We numerically integrated this system using ode45 in Matlab with a time step of 0.03. We visualized the delay space using data from $t = 0$ through $t = 29.97$ (time indices 1-1000). We used the delay vector parameters ($E = 3, \tau = 3.6$). For CCM with temporally separate training and testing sets, we used data from $t = 0$ through $t = 14.97$ (time indices 1-500) for training data and data from $t = 15$ through $t = 29.97$ (time indices 501-1000) for testing. For CCM with temporally interspersed training and testing sets, we used cross-validation over the entire range $t = 0$ through $t = 29.97$.

Third row: For this system, we used the initial conditions $W(0) = 0, \dot{W}(0) = 1/1.5, X(0) = 1.3, Y(0) = 1.5$. We numerically integrated this system using ode45 in Matlab with a time step of 0.1. We visualized the delay space using data from $t = 0$ through $t = 40$ (time indices 1-401). We used the delay vector parameters ($E = 3, \tau = 3.0$). For CCM with temporally separate training and testing sets, we used data from $t = 0$ through $t = 19.9$ (time indices 1-200) for training data and data from $t = 20$ through $t = 39.9$ (time indices 201-400) for testing. For CCM with temporally interspersed training and testing sets, we used cross-validation over the entire range $t = 0$ through $t = 39.9$.

Bottom row: We discretized this system with a time step of 0.05. We visualized the delay space using data from $t = 0$ through $t = 20$ (time indices 1-401). We used the delay vector parameters ($E = 2, \tau = 2.5$). For CCM with temporally separate training and testing sets, we used data from $t = 0$ through $t = 9.95$ (time indices 1-200) for training data and data from $t = 10$ through $t = 19.95$ (time indices 201-400) for testing. For CCM with temporally interspersed training and testing sets, we used cross-validation over the entire range $t = 0$ through $t = 19.95$.

For convergent cross mapping, we used the same $\tau$ and $E$ as for visualizing delay spaces (see above). "Training data size" on the horizontal axis is the number of delay vectors in the training library. Each dot in these CCM plots represents the average forecast skill over 300 randomly chosen libraries. Error bars represent the 95% confidence interval as calculated by the bias-corrected and accelerated bootstrap (1000 bootstraps) as implemented in Matlab's bootci function. Error bars are the same color as the dots and so are not visible when they fit inside the dots.

#### Methodological details for Figure S16

Top row: For this system, we used the initial conditions $X(1) = 0.2, Y(1) = 0.4$ and composed delay vectors of length $E = 2$ with a delay of $\tau = 2$. We visualized the delay space using data from time points 501-2000. We used points 801-1000 for training data and points 1001-2000 for testing cross map predictions.

Second row: For this system, we used the initial conditions $X(1) = 0.4, Y(1) = 0.2$ and the delay vector parameters $(E = 2, \tau = 1)$. We visualized the delay space using data from time points 501-2000. We used points 801-1000 for training data and points 1001-2000 for testing cross map predictions.

Third row: For this system, we used the initial conditions $X(1) = 0.2, Y(1) = 0.4$ and the delay vector parameters $(E = 3, \tau = 2)$. We visualized the delay space using data from time points 501-2000 (time points 1-6 $\times$ $10^5$ for the zoomed-in inset). We used points 801-1000 for training data and points 1001-2000 for testing cross map predictions.

Fourth row: For this system, we used the initial conditions $W(0) = Y(0) = 0$ and $X(0) = \dot{W}(0) = \dot{Y}(0) = 1$. We numerically integrated this system using ode45 in Matlab with a time step of 0.1. We visualized the delay space using data from $t = 50.1$ through $t = 200$ (time indices 501-2000). We used the delay vector parameters $(E = 3, \tau = 7.2)$. We used data from $t = 70.1$ through $t = 100$ (time indices 701-1000) for training data and data from $t = 100.1$ through $t = 200$ (time indices 1001-2000) for testing cross map predictions.

Fifth row: For this system, we used the initial conditions $X(1) = 0.2, Y(1) = 0$ and composed delay vectors of length $E = 2$ with a delay of $\tau = 2$. We visualized the delay space using data from time points 501-2000. We used points 801-1000 for training data and points 1001-2000 for testing cross map predictions.

Sixth row: For this system, the "initial" conditions specified the first 3 timepoints since we included a lag of 3. Thus, for $k = 1, 2, 3$, $W(k) = 0.2$, $X(k) = 0.4$, and $Y(k) = 0.3$. We composed delay vectors of length $E = 3$ with a delay of $\tau = 1$. We visualized the delay space using data from time points 501-2000. We used points 801-1000 for training data and points 1001-2000 for testing cross map predictions.

For convergent cross mapping, we used the same $\tau$ and $E$ as for visualizing delay spaces. The training data size is the number of delay vectors in the training library. For the plots in the fourth column, we chose 300 random libraries of training delay vectors with variable training data size, and used the standard prediction lag of 0. Delay vectors were chosen without replacement. Note that at large training data size, some or all of the 300 random libraries can be identical. Each dot in these CCM plots represents the average forecast skill over all 300 randomly-chosen libraries. Error bars represent the 95% confidence interval as calculated by the bias-corrected and accelerated bootstrap (1000 bootstraps) as implemented in Matlab's bootci function. Error bars are the same color as the dots and so are not visible when they fit inside the dots.

In all rows, the cross map skill for the putative causer $Y$ was greater than for at least 95% of random phase surrogate time series (purple dot). The 5% cutoff value was computed for the maximum library size (156 for row 4 and $\sim 200$ for all other rows) by running the CCM procedure after replacing the putative causer $Y$ with 500 random phase surrogate time series generated using the rEDM function make_surrogate_data.

For the plots in the fifth column we used the full library contained within the training data window (156 delay vectors for row 4 and $\sim 200$ for all other rows) and varied the prediction lag. We did not use random libraries for these plots.

#### Methodological details for Figure S17

To generate randomized parameter sets, we randomly selected $R_X$, $R_Y$, $A_{XY}$ and $A_{YX}$ from uniform distributions. We also randomly selected the initial conditions $X(1)$ and $Y(1)$ from uniform distributions. To make systems in the "friendly" parameter regime, we drew $R_X$ and $R_Y$ independently from the range $3.7 - 3.9$, we drew $A_{XY}$ and $A_{YX}$ independently from the range $0.05 - 0.1$, and we drew $X(1)$ and $Y(1)$ independently from the range $0.01 - 0.99$. These are the same parameters used in the randomized numerical simulations of [3]. Next, to make systems in the "pathological" parameter regime, we drew $R_X$ from the range $3.7 - 3.9$, we drew $R_Y$ from the range $3.1 - 3.3$, we drew $A_{XY}$ and $A_{YX}$ independently from the range $0.15 - 0.2$, and we drew $X(1)$ and $Y(1)$ independently from the range $0.01 - 0.99$. For both parameter regimes we randomly chose 500 sets of parameters and ran the system for 3000 time points. Occasionally a randomly chosen system would leave the basin of attraction and reach large values, represented on the computer as positive or negative infinity, or "not a number". When this occurred, we discarded the data and resampled parameters.

To apply CCM on each system, we generated a training library of delay vectors of $X$ by randomly selecting 200 vectors from among time points 100-2000. We then evaluated cross map skill from delay vectors of $X$ to values of $Y$ at points 2001-3000. Following [3], we used delay vectors of length $E = 2$ and a delay duration of $\tau = 1$. We evaluated cross map skill with a prediction horizon of $-8$ through 8.
